# Who’s driving? Common evolutionary mechanism of activation of class A GPCRs

**DOI:** 10.64898/2026.06.25.734477

**Authors:** Antoni Marciniak, Paweł Kozielewicz, Darko Mitrovic, Lucie Delemotte

## Abstract

Cells communicate with their environment by integrating signals, often chemical in nature, triggered by specific molecules bind to specific membrane-bound receptors, resulting in a downstream signaling cascade. Arguably, Gprotein-coupled receptors (GPCRs) constitute the most pharmacologically important family of such receptors, binding small molecules, peptides, lipids, and hormones with high specificity. However, despite a highly conserved fold and sequence similarity, GPCRs are still mostly studied on a case-by-case basis. Here, we infer a general, evolutionarily conserved mechanism of class A GPCR activation. By leveraging coevolution and machine learning methods applied to all class A GPCRs structures, we derive a mathematical description (a so-called collective variable - CV) of the receptor’s activation state which is independent of its sequence. Then, we bias molecular dynamics simulations along this CV to obtain transitions between activation states of a diverse set of class A GPCR family members. To demonstrate that our model generalizes beyond GPCRs in our training set, we obtain conformational transitions of an orphan receptor, GPR183. Finally, we show that we can model ligand effect on the receptors by converging Free Energy Surfaces of activation of the *β*2-adrenergic receptor within this common mechanism framework. These results, to our knowledge, prove for the first time the existence of a mechanism uniting all class A GPCRs. Our approach thus facilitates direct comparisons between receptors and opens up the possibility of structural and dynamical studies of many orphan and understudied GPCRs. It also serves as a blueprint for inferring family-wide protein mechanisms.

## Introduction

G-protein coupled receptors (GPCRs) are the largest class of membrane proteins in humans. They bind ligands with diverse chemical structures, and modulate a variety of physiological functions, ranging from the fight or flight response, metabolism regulation, immune reactions, to muscle-to-brain signaling.^1^ Despite the same fold and high sequence similarity, they respond to different stimuli and exhibit a limited propensity to cross-bind orthosteric ligands. These properties make them exceptionally attractive drug targets: over 36% drugs on the market target GPCRs.^2^ However, this high degree of similarity makes their exploration an incredibly hard task: with only a single common major motion TM6 outward movement - differences between receptors come down to minor structural changes that fine-tune the speci-ficity. These minor structural changes usually involve side chain rearrangements, which are often so subtle that they are impossible to capture with modern structural biology techniques. In fact, even capturing stable conformations poses a significant challenge: most of the GPCR structures have been determined using factors stabilizing them in a given state, such as ligands, binding partners (G-protein or arrestins), stabilizing inserts, bound nanobodies, sets of mutations or even as a chimera with an entire helix from another receptor.^3,4^ In addition, many studies based on spectroscopy techniques such as NMR, DEER, BRET and FRET suggest highly dynamic conformational landscapes.^5–8^ Thus, while available structures have facilitated a great deal of scientific and pharmacological developments, it remains unclear to what extent they reflect the thermodynamics of receptors in physiological conditions. In addition, the description of the conformational state captured in structures is often based on so-called “microswitches”, particular side chain rearrangements of well-conserved amino acids recognized in previously-solved structures of, possibly, other receptors. This risks propagating bias from structure to structure. In fact, many of the receptors do not preserve putatively characteristic amino acids, making direct comparisons challenging (Table S1). In addition, an issue with microswitch-based state description is that it remains unclear if these microswitches actually energetically drive the conformational change, or are they just “passengers”, i.e. highly correlated proxies for yetto-be identified drivers that truly define the state of the receptor.^9^ As a consequence of this, systematic comparative studies of GPCRs remain scarce despite the preservation of the fold and downstream signaling partners. Nevertheless, the specialization of receptors implies the existence of a common mechanism that could theoretically be traced to an ancestral, proto-GPCR a mechanism that is present today in modified forms across GPCRs. Attempts to find a common frame of reference between different receptors started in 1995 with the seminal study which resulted in the widely used Ballesteros-Weinstein numbering. ^10^ More recently, several studies have focused on unifying the microswitches’ patterns within different receptor classes by comparing published structures.^11,12^

In this work, we sought to find a common, unifying mechanism of activation for all class A GPCRs. To achieve this, we analyzed evolutionarily important distances in all available structures and pinpointed contacts that universally change between active and inactive states. We then built a collective variable based on a selection of the aforementioned distances and biased the sampling of molecular dynamics simulations along these CVs to nudge receptors to transition from the active to inactive state and *vice versa*. We then showed that this procedure is transferable to an orphan GPCR, GPR183, which was not represented in our training set. By projecting all of our transition data to a common 2D space which connects active and inactive states, we could compare the activation mechanism across the class A GPCR superfamily. Finally, we showed that our description of activation can be used to converge Free Energy Surfaces of the activation of the *β*2-adrenergic receptor activation in apo form and in presence of ligands of various modalities, serving as a proof of concept that our pipeline can be used to compute the free energy landscape of activation of any class A receptor and evaluate how ligands affect the activation.

## Results

### The class A GPCR activation mechanism is evolutionarily conserved

To start off, we set out to define a mathematical description of class A GPCR conformational state, commonly called a collective variable (CV) in the field of molecular modeling. We built a dataset containing 447 structures of class A GPCRs (188 in the active state and 259 in the inactive state, extracted from GPCRDB in 2022-11) and chose to build this CV based on distances between residue pairs. Indeed, previous work has highlighted that most confor-mational changes are driven by local changes in contacts between amino acids, making a mathematical description focused on distances that describe contacts that are formed or broken during the conformational change generally powerful. ^13–16^ In addition, descriptions based on inter-residue distances have the advantage of being rotation- and translation-invariant, removing the need for structural alignments that can be ambiguous.^17–20^ To select only the distances that are important for all class A GPCR activation mechanisms, we focused on highly co-evolving residue pairs, or on pairs of highly conserved residues. Coevolution is a marker of structural proximity: indeed, pairs of residues which need to be in spatial proximity to facilitate various aspects of protein function, be it folding, binding another protein or stabilizing a conformation, will tend to coevolve. Obtaining relative coevolution scores (couplings) can be done by fitting a Pott’s model using a multiple sequence alignment (MSA),^21^ and the aforementioned couplings have been used to predict protein structure, find interacting proteins, find new conformations and sample protein conformational landscapes, among others.^13,14,16,21–29^ On the other hand, conserved aminoacids usually do not show up in highly coevolving pairs since there is little sequence variation to correlate with changes elsewhere in the sequence. Thus, to also pick up potentially functionally important distances between conserved aminoacids, we created all possible combinations of pairs of highly conserved residues, defined as positions where the frequency of a single amino acid exceeds 50 percent. This procedure yielded a set of 594 potentially evolutionarily important pairs of residues (Fig1.A).

**Figure 1.**
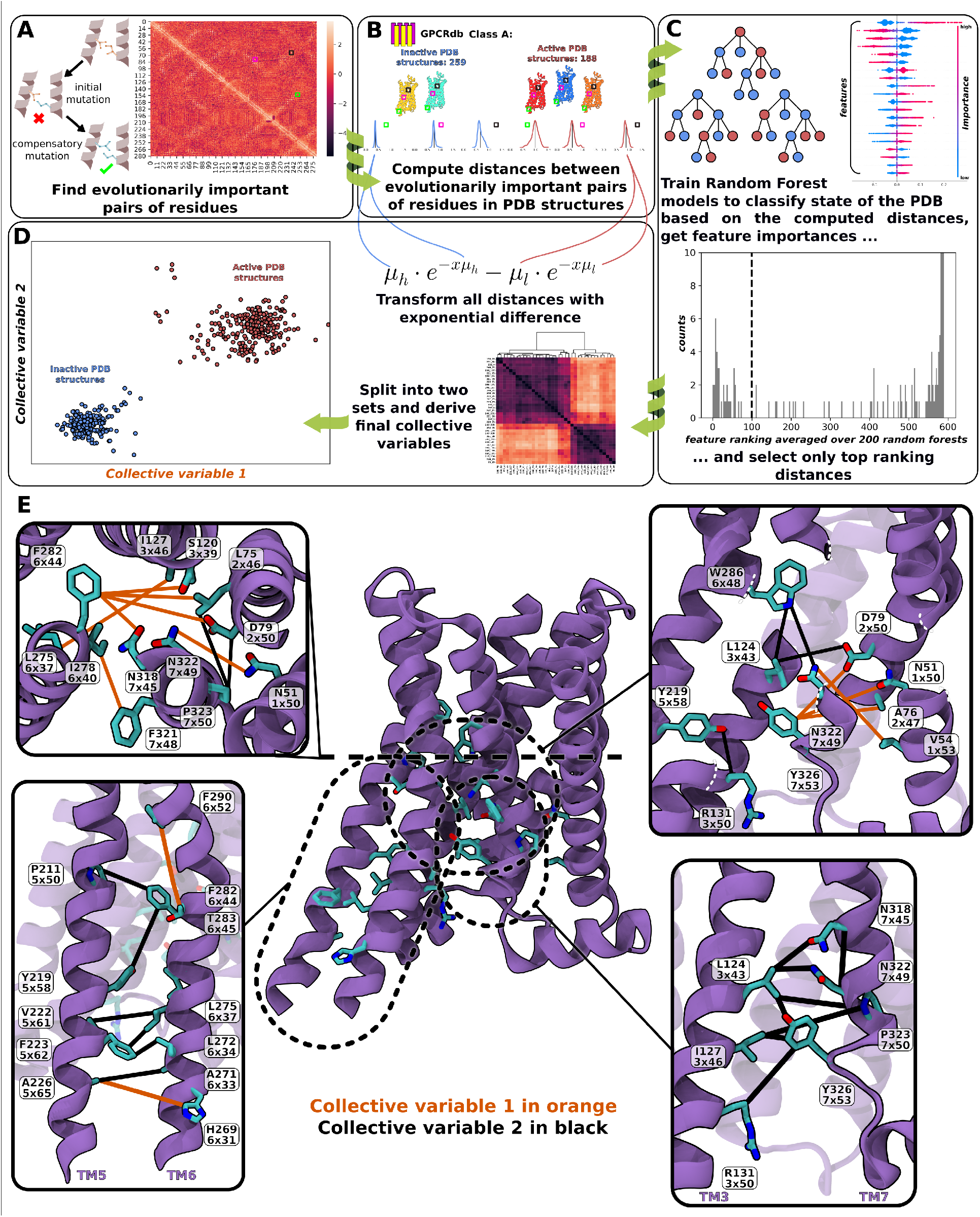
Blueprint for deriving (co)evolution-based collective variables(CVs). First, we computed coevolutionary couplings and picked only ones above 2 standard deviations from the mean, as well as all possible pairs of conserved residues defined as at least 50% frequency of a single amino acid at a particular position in the Multiple Sequence Alignment (MSA) (panel A). We then computed distances between these pairs in all structures of class A GPCRs listed on GPCRdb in 11.2022 (panel B). Next we trained 1000 separate random forest classifiers with these distances to classify the state of the structure as active or inactive, computed feature importance with Shapley values, ranked them for each model, computed an average rank for each pair, and selected the highest ranking ones (panel C). For those, we split them further into two sets by clustering them based on their linear correlation, and transformed each of them with difference of exponentials where *µ* parameters are mean values of the distance in the pdb structures (panels B&D). Finally, we used Harmonic Linear Discriminant Analysis (HLDA) on both sets independently of each other to obtain linear coefficients for the final expressions for collective variables (panel D). Projections of active PDB structures on the 2D HLDA expression are shown in red while projections of inactive PDB structures are shown in blue. Physical representations of the residue pairs used in the CVs are shown in panel E, with the line colors corresponding to the colors of CVs as shown on the panel D (orange for CV1 and black for CV2). Residues are shown on the active *β*2-adrenergic receptor structure (PDBID 4LDO), with matching Ballesteros-Weinstein (BW) numbering below. Our residue numbering is based on our MSA.

As mentioned before, coevolution may reflect spatial proximity at any point of the protein’s life cycle. To focus on the pairs involved in the conformational change related to activation, we sought to further filter potentially important pairs by systematically comparing active and inactive structures. To do so, we used explainable machine learning to extract the pairs that best separate active and inactive states. We first computed distances be-tween all 594 evolutionarily important pairs in all class A structures (Fig1.B). Next, we trained 200 Random Forest models to differentiate between active and inactive PDB structures using the aforementioned distances as features, and ranked features from each model according to their feature importance, as computed as Shapley values.^30,31^ Then, for each of the features, we computed its average rank across all 200 models, and selected only the highest ranking features as potential drivers of the conformational change. We picked Random Forest as we wanted to use ensemble learning applied to non-linear models with, *i*.*e*. leverage a set of models of the same category that collectively “vote” on the final outcome, thus lowering the model’s variance (Fig 1.C).

### Our collective variable can drive the conformational change of a broad range of receptors

Next, we wanted to test if the distances we uncovered could be considered drivers of the conformational change associated with activation. To achieve this, we turned to molecular dynamics, as it is currently the only technique that is capable of describing conformational dynamics in atomistic details, and sought to define a collective variable based on these distances that would allow us to bias the dynamics and observe transitions between activation states. One of major pitfalls when designing CVs is degeneracy. The more degenerate a collective variable is, the least efficient it is at sampling the process of interest. Therefore, we first split our sub-selected distances into two sets based on their linear correlation. Such split reduced degeneracy, and possibly helped separate directions of motion of receptor parts. Then, we used Harmonic Linear Discriminant Analysis applied separately to the distances contained in each of two final splits, obtaining a 2D description of the receptor state (Fig.1.D).^32^ As our distances were defined through positions in the MSA, a particular pair may involve different amino acids in different receptors such that, at a given distance, residues may or may not be in contact based on their physicochemical properties. To compensate for such differences, we transformed each distance into difference of negative exponentials (Fig.1.D). This way, distances with small and large ranges are treated with similar importance and shorter distances are prioritized.

Having obtained a mathematical description of receptor structure, we could test if we captured causal factors behind conformational change by directly biasing the sampling of MD simulations along this collective variable set (Fig.2.A). To make these tests robust within the class A family, we picked 20 different receptors within 20 different class A subfamilies (Fig. S1, Table S2). We looked for GPCRs with structures available in both active and inactive states, and picked receptors to cover a wide range of sequences. Notably, this collection of receptors contains exceptions with respect to the most commonly considered microswitches: three lack tryptophan (6x48) in the CWxP motif (Orexin receptor 2, Prostaglandin E2 Receptor EP4 subtype and Thyroid Stimulating Hormone Receptor), two lack tyrosine (5x53) in the tyrosine lock (Prostaglandin E2 Receptor EP4 subtype and Neurotensin receptor type 1); one lacks the asparagine (7x49) in the NPxxY motif (Prostaglandin E2 Receptor EP4 subtype); four lack the conserved serine (3x39) in sodium pocket (Prostaglandin E2 Receptor EP4 subtype, Rhodopsin, Neurotensin receptor 1, and ghrelin); and five lack the asparagine (7x45) in the sodium binding pocket (Chemokine, Rhodopsin, Neurotensin, Ghrelin, Neuropeptide Y) (Fig.2.B, Table S1). We let the structures relax to their thermodynamic minima by running short equilibrium simulations of all 40 structures: 20 receptors in active and inactive states. Then, we biased sampling using the accelerated weight histogram (AWH) method (Fig.2.A). To ensure further robustness, we started transitions from the active state for 12 of the receptors and from the inactive state for the other 8 (see SI Receptor Data). We verified that transitions were completed (i.e. reached the other state) by 1-computing C*α* root mean square deviations (RMSD) between transition trajectories and equilibrium ensem-bles, 2-computing popular microswitch conformations along the trajectories, and 3-inspecting visually all transition trajectories. Moreover, we confirmed that applied bias did not induce breaking of secondary structure elements as shown by the secondary structure evolution over time (Fig. S2-S61). Encouragingly, all receptors transitioned successfully within 1 *µ*s. As expected, the time it took to reach the other state varied between receptors. Note that this time does not generally correlate with the receptors’ kinetics of activation given that our CV does not consider the sequence specificity of each receptor and is not expected to account for all the slow degrees of freedom related to activation of a given receptor.

**Figure 2.**
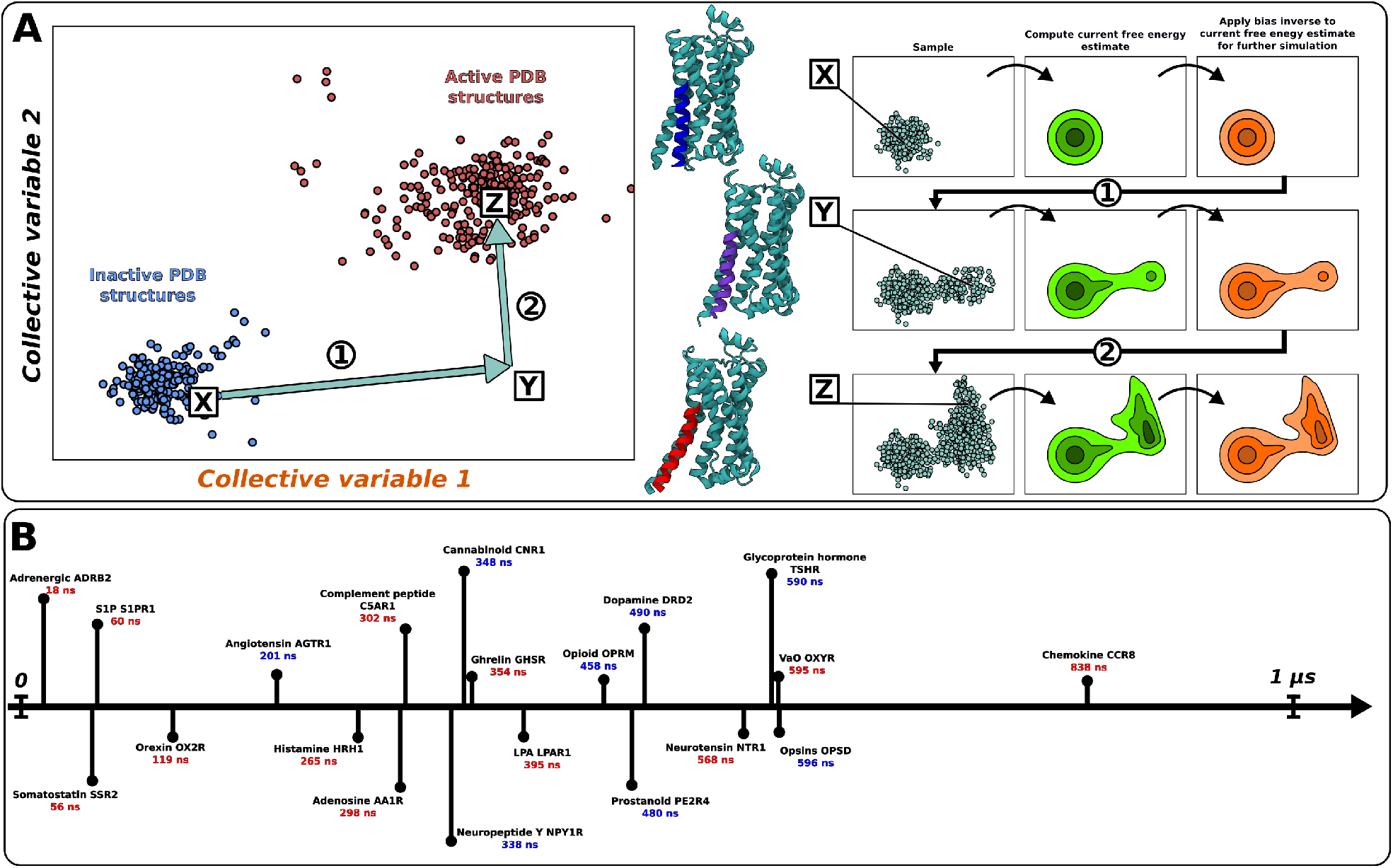
(A) Adaptive biasing algorithms (such as AWH) explore conformational space very efficiently. In this procedure, first, unbiased simulations are run. Next, based on the collected samples, a current free energy estimate is computed and its inverse is applied as a bias. The simulation is then continued with this bias applied and new samples are collected again. Next, the free energy is estimated using all samples collected until that moment, and its inverse is again applied as a bias. This procedure repeats until convergence. Sampling within our CV space has been shown schematically, with structures at position X signifying inactive states, at position Y signifying intermediate states, and as position Z signifying active states. ^33–35^ (B) All 20 receptors whose sampling is biased along our CV transition to the other conformational state. Timeline of simulation times necessary for the first transition to the opposite state for all 20 class A receptors. We considered that a receptor has transitioned when RMSD traces crossed over, and TM3-TM6 distance crossed the average value for the ensemble in the opposite state (SI appendix Fig. S2-S61, Fig. 4). For each sequence, the receptor, gene name and time until first transition are given. We started transitions for receptors with transition times in red in the active state and in blue in the inactive one.

### A common mechanism connects activation states even in high dimension

As only selected distances were included in the CVs, we wanted to ensure that transitions also connected both states if we took a broader perspective, e.g. include all the distances that highly coevolve and all pairs between conserved residues. To this end, we trained an autoencoder on the short equilibrium simulation trajectories of the 20 representative receptors in the active and inactive states, using distances between all 594 evolutionarily important pairs as features (Fig. 3.A). We then projected the transition trajectories to the 2D latent space of the autoencoder. This way, we could visualize all simulations in a common, non-linear space. As can be seen on Fig. 3, most of the active/inactive ensembles are connected by their transitions. Our projection also clearly separates active and inactive ensembles, and many receptors overlap with their closest phylogenetic relatives. We consider the common orientation and direction of projections of transition trajectories a further proof that we have captured a nonlinear manifold describing the activation mechanism common to various class A GPCRs. While the common 2D autoencoder projection further highlights that transitions occur even when considering many more distances than those considered in the CV used for biasing, the equilibrium ensembles of a few receptors do not appear perfectly connected by the transition data: we believe that slight disconnections between transition and equilibrium ensembles occur due to the fact that the autoen-coder is sequence-unaware: as mentioned earlier, a particular pair may be involve different amino acids in different receptors, potentially leading to multimodal distributions of features where minor modes are likely dampened by the major modes, leading to disconnections in the low-dimensional projection. Altogether, we fur-ther confirm that our transitions connect both states even in higher dimensions.

**Figure 3.**
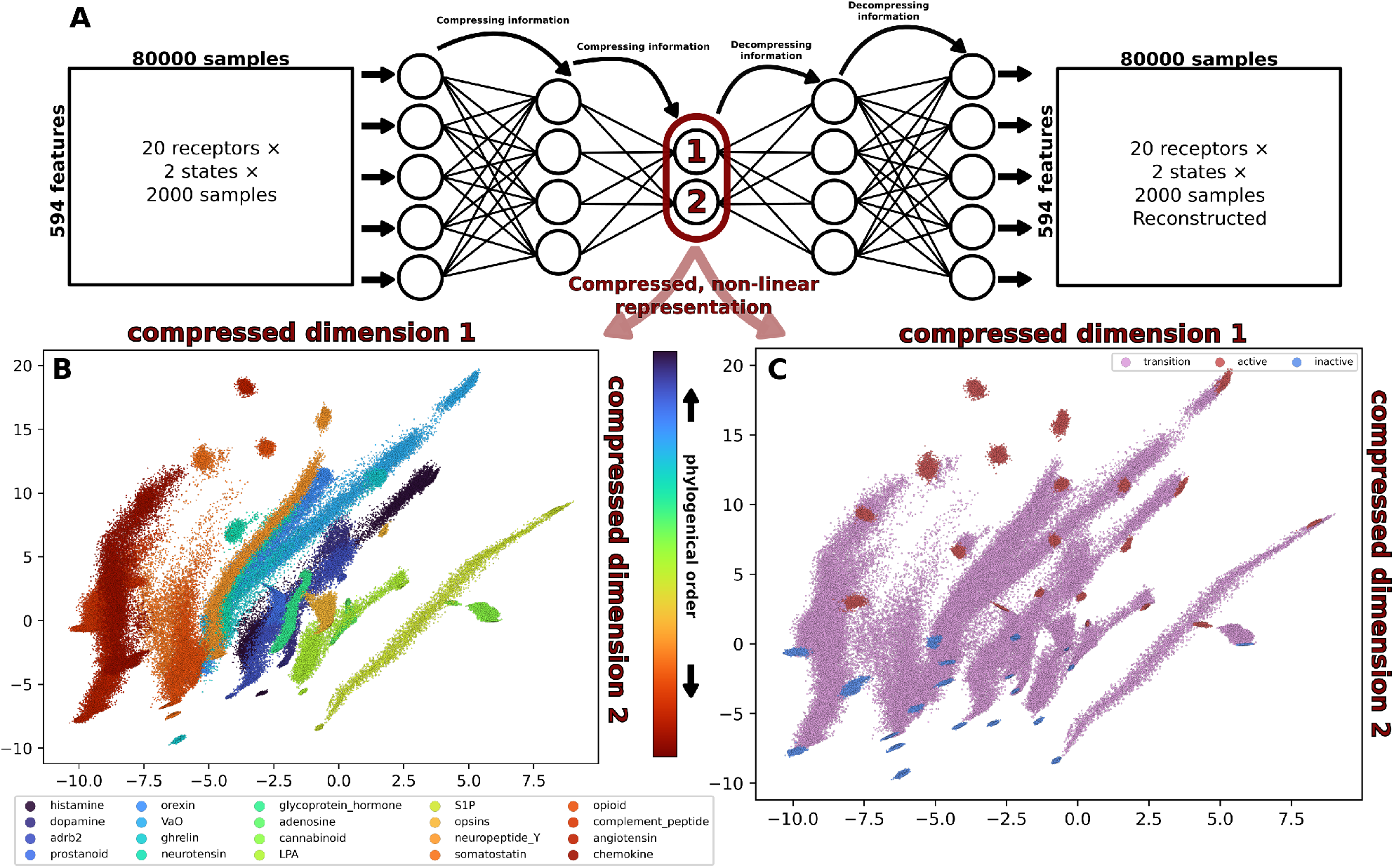
A common 2D projection clearly separates active and inactive ensembles, transitions connect active and inactive ensembles, and many receptors overlap with their closest phylogenetic relatives. (A) Schematic representation of an autoencoder. An autoencoder is a deep learning model in which the training goal is to reconstruct the input data. The size of the layers progressively decreases until the smallest-sized layer in the encoder part of the model, and then size of the layers progressively grows in the exact reverse in the decoder part of the model. This way, the model learns to compress the information and stores the most information-efficient representation in its smallest layer (the bottleneck, also called the latent space). To train our model, we have used all 594 evolutionarily important distances computed from active and inactive equilibrium simulations of 20 receptors. We then projected the transition trajectories into its bottleneck layer. Projections of MD data onto the 2D latent space. In (B), we have colored the projections by receptor sequences, ordered according to their phylogenetic proximity (Fig. S1). In (C), we have colored the projections according to their state: equilibrium trajectories of receptors in active and inactive states are shown in red and blue, respectively, and transition trajectories in purple. For clarity, every other simulation frame has been projected.

### Our general CV can drive the conformational change of an understudied, out-of-training receptor

Having confirmed that our CVs are able to describe and drive class A GPCR activation across the 20 receptors part of the structural training set, we wanted to test whether they could be used to guide the transition of a receptor that had structure of a single state included in the original training set. We chose GPR183 as one of the few orphan receptors with structures of both states available, and purposefully excluded its inactive structure from our dataset, reserving it for validation purposes.^8,36^

We started from the active structure of GPR183 (PDB ID: 7TUY) and biased simulations along the family-wide CV described above to reach the inactive state. As shown by RMSD traces (Fig. 4), for this recep-tor too, we quickly reached the inactive state while maintaining the secondary structure (Fig. S64) intact and sampling a broad range of microswitch conformations (Fig. 4D and Fig. S63). Notably, GPR183 has exceptions to known microswitches: Phe in the place of Trp in CWxP motif, and Asp in the place of N in NPxxY. Projection of both equilibrium and transition simulations to the latent space of the autoencoder mentioned above also shows that GPR183 embeds itself as expected, with the active and inactive conformational ensembles projecting in the same area than the active and inactive states of the other GPCRs, respectively, and the transition trajectory connecting both equilibrium ensembles and mapping closest to phylogenetically-related sequences (Fig. 4). These results further solidify our CVs as general drivers of Class A GPCR conformational changes.

**Figure 4.**
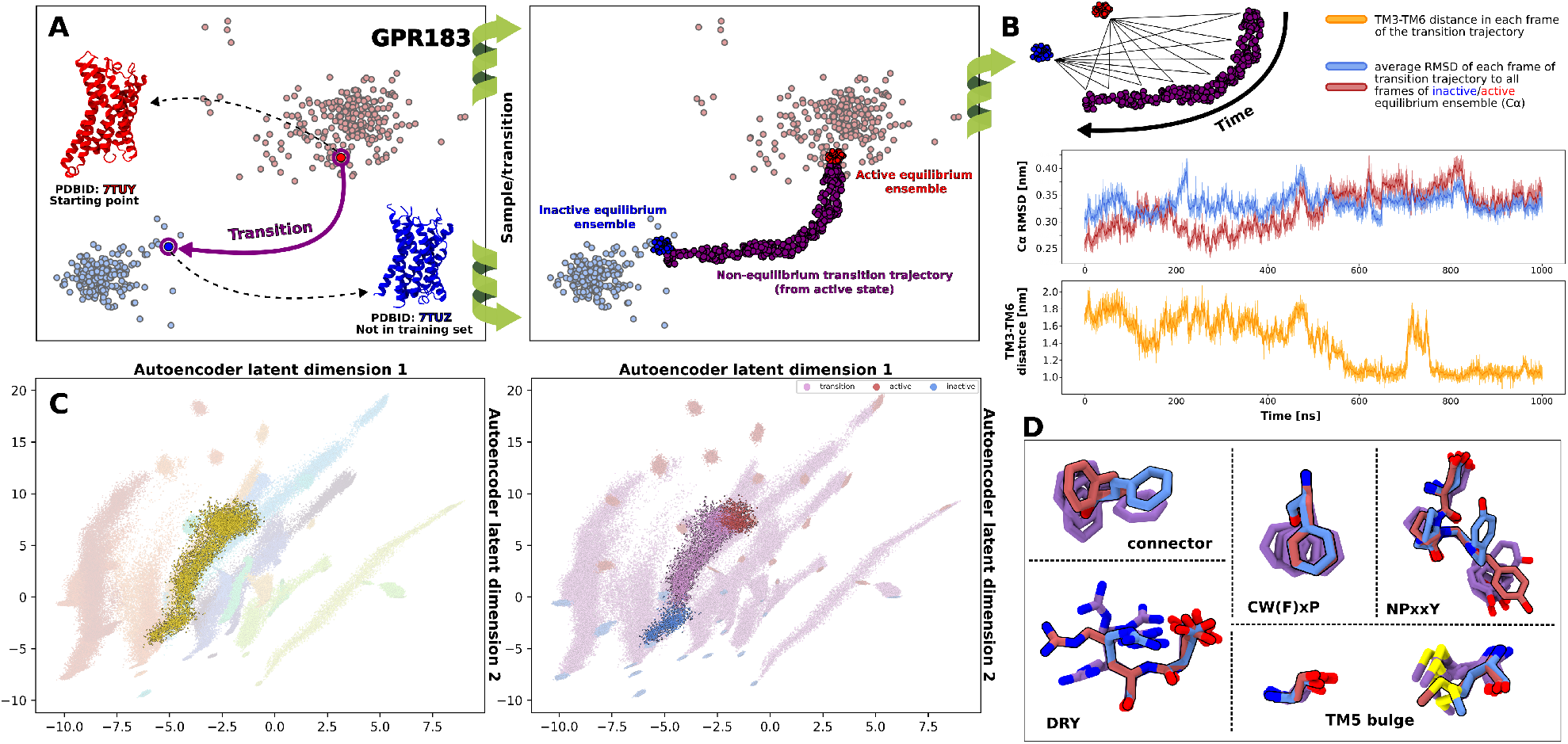
Our pipeline can also guide the activation transition of the orphan receptor GPR183. (A) We started transition simulations from active structure (PDBID: 7TUY), and reached the inactive state (PDBID: 7TUZ, purposefully withheld from the training set). (B) To validate the transition, we computed a transmembrane C*α* RMSD from the active and inactive ensemble along time, in red and blue, respectively, and the TM3-TM6 distances (measured as the distance between the C*α* of residues 6x32 and 3x50) along time. On the both plots, per frame values are shown in lighter shades and running averages taken over 25 frames are shown in darker shades. (C) A projection of GPR183 simulations to the autoencoder latent space is shown. For clarity, only every second point has been projected. (D) Snapshots of classical microswitches from the transition trajectory, selected to showcase the range of states that were sampled. As GPR183 has Phe in place of Trp in the CWxP motif, we visualized Phe motions as representative of this motif.

### Various ML models highlight similar inter-residue distances as important for activation

To verify if a much more powerful model, e.g. an autoencoder, focuses on similar distances, we sought to understand which features are the most important for establishing its latent space. To unravel this, we performed layer-wise relevance propagation analysis (LRP) on the data from different receptors^37^ (Fig. 5 and SI appendix, Fig. S65-S68). In agreement with our expectations, almost 50% of distances establishing both latent dimensions are present in our CVs (when considering distances important for at least 15 out of 20 receptors). We hypothesize that distances absent from the CVs could be so-called passengers, distances that highly correlate with some of the pairs included in the CVs but contain no new information compared with the pairs that were included. Moreover, compared with the ensemble of 200 Random Forest models used to derive the CVs, we expect an autoencoder model to have much higher variance. As shown on the *β*2AR structure, many of important distances revealed by the AE analysis are located in or near extracellular loops. These distances would fluctuate a lot even in equilibrium MD simulations and contribute noise to the model. Interestingly, none of the pairs important only for a single receptor were included in the CVs (Fig. S67,S68), further cementing the idea that our CV captures general, common mechanism while omitting receptor-specific degrees of freedom. Overall, as the much more complex model predicts 12 of the same pairs to be important as the ensemble of Random Forests, our CV choice is further validated; in fact, if we have chosen only the features ranking in the top 20 on average in our CVs derivation, instead of the top 100, 9 pairs remain common with the AE.

**Figure 5.**
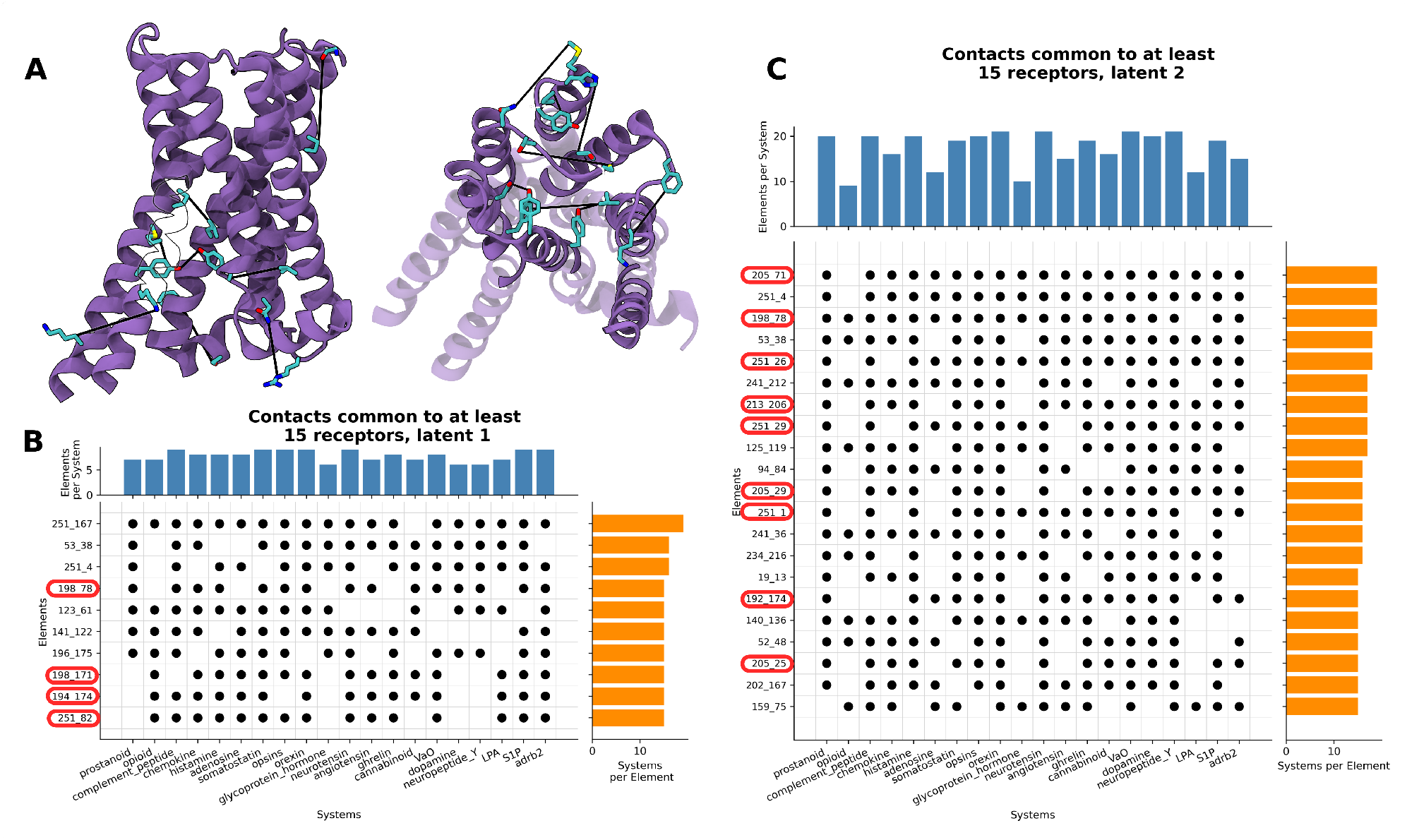
Upset plots of the most important features in latent dimension 1 (B) and latent dimension 2 (C), as revealed by Layerwise Relevance Propagation. For each of the 20 receptors in our training set, the top 100 most important distances were identified, and only ones present in the top 100 for at least 15 systems are shown. Histograms on the right count for how many of the 20 systems the distance was in top 100, whereas histograms on the top count how many of top 100 distances are present in each of the systems. We marked distances in our general CVs in red, and visualized the absent ones on the *β*2-adrenergic receptor structure on (A).

### Free energy surfaces converged using our CV can distinguish ligand modalities

Finally, to test if we could capture the effect of ligand binding, we computed the free energy surfaces (FES) of *β*2-adrenergic receptor activation with a variety of ligands with different downstream signaling effects: the beta-blockers alprenolol, timolol and carazolol; the partial agonist salmeterol; the orthosteric agonists adrenaline and the synthetic agonist formoterol, and in the apo condition. ^38^ Since we are now focusing on a single protein, we could further refine the general CV to speed up the convergence of the FES. We did so by performing Principal Component Analysis (PCA) on the transition trajectory of the *β*2-adrenergic receptor, using as input the distances included in the general CVs presented above. We then used the first two principal components as CVs (SI appendix). As shown on Fig. 6, the apo receptor clearly favors inactive states. When bound with any drug, the most stable states are either active or intermediate. While this is not the expected outcome for beta blockers, the simulations have been conducted in the presence of a protonated D2.50, a condition known to heavily favor activation.^39,40^ Therefore, we believe that our profiles are all shifted towards active states. Contrary to our expectations, it is the ratio between intermediate and inactive states that correlate with the ligand efficacy (Fig. S69,S70). That is in agreement with previous long scale MD studies, where Dror *et al* suggested that G Protein binds to *β*2-adrenergic receptor in an intermediate state. Furthermore, there are several works showcasing G Protein coupling to other receptors in inactive or intermediate state.^5,7,41–43^ It thus appears that beta blockers are the only drugs that do not stabilize intermediate states significantly. Our results therefore suggest that sta-bilization of intermediate states is necessary for agonist activity. Moreover, further stabilization of active states given stable intermediates correlates strongly with agonist efficacy (For state assignments, see Fig. S69, for convergence analysis see Fig. S71-S83).

**Figure 6.**
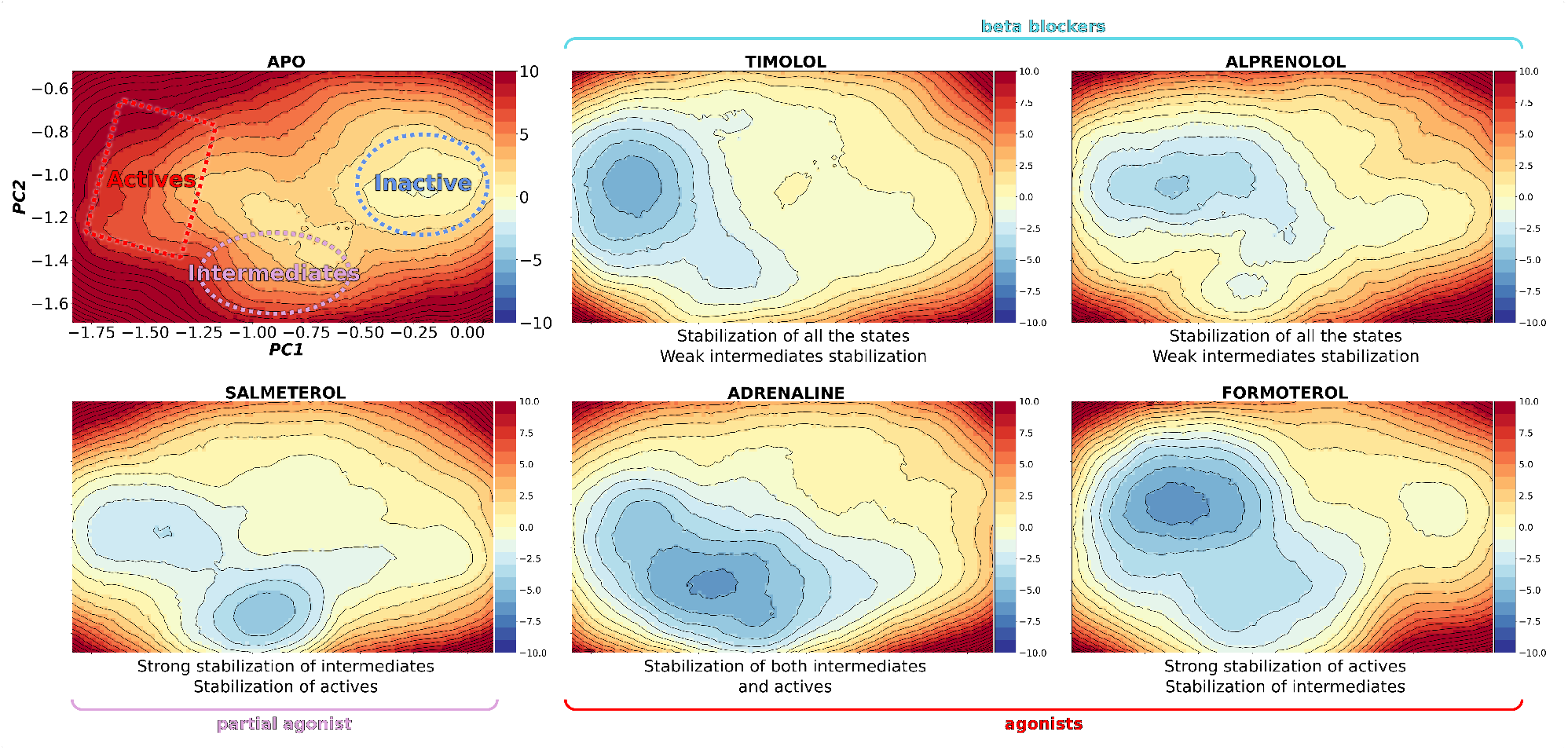
Free energy calculations predict ligand function. Free energy surfaces of the apo *β*2-adrenergic receptor in the apo condition (left top row), bound to beta blockers timolol and alprenolol (middle and right top), bound to partial agonist salmeterol (bottom left), bound to orthosteric agonist adrenaline (middle bottom) and bound to synthetic agonist formoterol (bottom right). All free energies are in kcal/mol, and the center of the inactive basin from apo condition has been chosen as reference point (with energy equal 0) for all surfaces.

## Discussion

We have uncovered a common activation mechanism relevant to a wide range of receptors devoid of the sequence-dependent issues present in classical microswitch analysis. Our description, based on coevolution and with available structures serving only as a filter, allows us to alleviate some of the biases stemming from structure determination studies. Our method facilitates access to conformational spaces of unknown and understudied receptors, such as the orphan receptor GPR183 that we have used for validation purposes, and only requires a single structure in any state as an input. Simulations of transitions presented in this work provide a plethora of intermediate, well-behaved configurations previously not seen in structural studies, thus widely broadening the known conformational space of GPCRs.

If we examine where residue pairs involved in our CV are located, it appears that the ligand binding site is completely devoid of them. There are, though, many contacts just beneath it, in what we call a spatial bottleneck: the region of the receptor where sodium binding site and 6x44 residue are located. Our work therefore aligns with previous findings by Dror *et al* : GPCR structures can be functionally divided into three parts: ligand binding site, downstream signalling domain, and a spatial bottleneck that couples them together. The coupling seems to work as a cascade from top to bottom: bound ligand shifts the conformational equilibrium of the binding site, which in turn shifts conformational equilibrium of the spatial bottleneck, which then shifts the conformational equilibrium of the downstream signalling domain.^41^

Why is our pipeline successful? We speculate that this is a result of a particular mix of modeling choices: first, narrowing down the most important features through nonlinear mathematical models and coevolution; second, CV formulation in which short distances are emphasized and long distances de-emphasized and forces depend on the current position in the CV space, i.e. as each distance included in the CV is transformed with exponential functions, derivatives of the applied potential (which will be applied as forces) are proportional to the po-tential itself, according to chain rule:

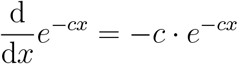

Third, the use of adaptive biasing algorithm which allows for a well-behaved application of the biasing potential. We believe that as long as these three elements are present, it should be possible to reproduce our results or apply the pipeline to a different family. Thus, this work also provides a blueprint for family-wide conformational exploration. In fact, this work was inspired by a simplified version of the pipeline we have used previously to obtain the whole conformational cycle of sugar transporters. ^14^

We would like to highlight that while a great body of work has been devoted to finding out key driving factors behind conformational change, to our knowledge, all analyze already transitioned trajectories retrospec-tively or static structures from the Protein Data Bank. In particular, Zhou et al analyzed all available class A GPCR structures using Residue Residue Contact Scores to narrow down 34 pairs of residues that rearrange upon activation. ^12^ The mechanism postulated in the aforementioned work was based on already known microswitches, and Zhou et al used it to design constitutively active/inactive mutants with a varying degree of success. Interestingly, 9 pairs identified by Zhou et al also came up in our analysis, mainly describing contacts between TM3 and TM6: 6x44 - 7x45, 3x46 - 6x37, 5x62 - 6x37, 3x50 - 7x53, 3x46 - 7x53, 3x43 - 7x53, 3x43 - 7x49, 7x45 - 7x49 and 2x50 - 7x49 (7x50 in our case). Moreover, 5 particular positions highlighted in their work serve as connection hubs in our CVs: 3 residues participate in 6 residue pairs: 6x44, 3x43 and 7x53; 1 in 5 residue pairs: 7x49, and 2 in 4 residue pairs 2x50 and 7x50 (not highlighted by Zhou et al). Mutations of these connection-hub residues were also reported in literature to have the strongest effect on the receptor activation. Later, Hauser et al pointed to 3x40, 3x46, 3x50, 5x58, 6x44, 7x53, 7x54 and 8x50 as residues that switch contact partner upon activation. 3x46, 3x50 and 5x58 appear in our CVs 3, 2 and 2 times, respectively, while 7x54 and 8x50 are not present at all (6x44 appears 6 times as mentioned above, and 3x39 instead of 3x40 appears once).^11^ In a seminal work by Heydenreich et al, authors tried to identify residues driving signaling in the *β*2-adrenergic receptor by mutating every single residue to alanine and testing its effect on adrenaline-driven signaling. Our CVs contain both residues classified as drivers (2x50, 3x39, 3x43, 3x46, 5x58, 6x37, 6x44, 6x52, 7x45, 7x49, 7x53, 39%) and passengers (1x53, 2x46, 3x50, 5x62, 6x40), 18%). However, it is hard to directly compare as this work reports downstream Gs signaling, whereas our CVs describe only the conformational changes of the receptor within the transmembrane region, in absence of G Protein. Moreover, while alanine mutations are often considered neutral, this is not always the case and such mutations can have major negative/positive impact on the local environment, thus altering the protein function instead of quantifying lack of specific amino acid at a certain position. ^9^ It is worth mentioning that all three aforementioned publications include contacts with helix 8, which was not present in multiple sequence alignment that our CV is based on. In fact, some of the structures we used in simulations lack helix 8 entirely, suggesting that it does not play a major role in the receptors’ conformational change, but remains crucial for interactions with binding partners. Finally, it seems that 3x46, 5x58, 6x44, and 7x53 emerge from all perspectives as absolutely crucial for *β*2-adrenergic receptor for all aspects of its function.

While there are many examples of successful molecular dynamics-based studies of GPCRs with enhanced sampling, most of them focused on a single receptor with only a few attempts to describe GPCR activation with more than a single sequence in mind. ^44–48^ However, even amongst these few, all analyses have been done retrospectively i.ex. analyzed already transitioned trajectories without testing causative factors; they also included a limited set of receptors, or necessitated structures of both states.^46,47^ Overall, our CVs build upon previous GPCR mechanism descriptions, with evolutionarily justified choice of contacts and wide receptor selection. Our CVs include regions with known microswitches while being independent of sequence, which we demonstrated by transitioning receptors with known exceptions to microswitches. Moreover, we used biased molecular dynamics directly to verify causality, as all receptors transitioned in sub-*µ*s simulation times, thus confirming that we capture a significant portion of causality. The fact that we were able to predict modalities of *β*2-adrenergic receptor ligands by computing free energy surfaces seems to further confirm that we captured a core part of mechanism. We did keep our CV for converging *β*2-adrenergic receptor FES simple on purpose to showcase the accuracy of using our pipeline out of the box: picking a receptor, obtaining a transition and then converging the FES in the first two principal components of the transition trajectories (SI appendix). Such a pipeline can be applied to any class A GPCR. However, this CV could be easily improved: we expect CVs derived through supervised learning (e.g. Support Vector Machines, Linear Discriminant Analysis) after clustering the transition trajectory to be much more detailed and accurate but this process requires individual optimization step for each receptor. Moreover, we also expect improvements should we add receptor-specific degrees of freedom. We would like to highlight that the only data used for obtaining the CV was a single transition of an apo receptor. Therefore, obtaining transition trajectories with ligands (or any other perturbation,e.g. mutation) would likely improve the accuracy without individual optimization to particular receptor, but at an increased computational cost. Using only an apo transition trajectory implies that it would also be possible to apply this pipeline to receptors that don’t have determined structures and can be only predicted by e.g. deep learning methods, especially given that our pipeline is starting state-independent. As described in the Supplementary Information, preparation of ligand-bound system was similar to docking, i. ex. initial ligand poses came from structures of the same receptor but were all copy-pasted to a particular structure. This highlights the possibility of using our pipeline to study compounds that have unknown binding pose, e.g. further evaluating top hits of docking screens. We hope to cover all of aforementioned improvements in a future work.

To sum up, our results prove that our CVs combined with physics-based modelling allow for a precise description of the activation process of all class A GPCRs, and can faithfully reproduce the influence of external factors on activation. As our pipeline is not inherently focused on GPCRs, we expect it to be applicable to any protein family, and any type of structural data could be used as a filter both of which will be a focus of our future work. Importantly, GPCRs physiological function depends on many factors, such as pH, lipid modulation and, most importantly, binding partners such as G-proteins or arrestins. This work does not address any of those interactions in detail, but provides a foundation to further test influence of these factors case-by-case with robust description of activation. Particularily, protonation of 2x50 is known to affect the conformational equilibrium of class A GPCRs.^39,40^ Many receptors are also known to bind cholesterol, and membrane tension is known to alter receptors’ conformational equilibria. All of our simulations were conducted in pure 1-palmitoyl-2-oleoyl-sn-glycero3-phosphocholine(POPC) membranes, and we hope to address both of these factors in a future study.

## Acknowledgement

We would like to thank Claudia Alleva, Marta Bonaccorsi, Jens Carlsson, Berk Hess, Samuel Eriksson Lidbrik, Jack Lindmar, Magnus Lundborg, Lukas Müllender, Simon Olsson, Adrien Schahl, Mi-losz Wieczór for useful discussions. This work was supported by the Knut and Alice Wallenberg Foundation (2019.0130), the Science for Life Laboratory, and the Swedish Research Council (VR 2019–02433 and 2022–04305). The National Academic Infrastructure for Supercomputing in Sweden (NAISS) and Swedish Research Council through grant agreement no. 2022–06725 funded MD simulations.

