## Supplementary material for "Who’s driving? Common evolutionary mechanism of activation of class A GPCRs": Tables S1-S3, Supplementary methods, Figures S1-S84

### 1 Methods

#### 1.1 Data curation

To obtain the dataset for collective variable design, we downloaded all class A GPCR structures from GPCRdb[1] (access: 11.2022). We excluded all structures with resolution lower than 4.5 Å and state assigned as 'Other' or 'Intermediate' by GPCRdb[1]. Next, we computed distances between all highly coevolving pairs (defined below) as well as between all combinations of highly conserved residues (defined as a position having conservation of a single amino above 50%). Out of these distances, we kept only the ones that were present in at least 95% of selected structures and were below 0.451 nm in at least one instance.

#### 1.2 CV derivation

To further subselect distances for CV design, we trained 200 Random Forest Classifier models to classify the states of structures - active and inactive, as labeled in GPCRdb[1] - based on the distances selected as described in "Data Curation" section. We trained Random Forests with 100 trees each, maximum depth set to 20, minimum samples per leaf set to 5 and minimum samples per split set ot 10%, as implemented in scikit-learn ensemble module. For each Random Forest, we computed Shapley values[2] for all the features (interresidue distances) as implemented in TreeExplainer in SHAP library[3], and ranked them. We then averaged the ranks across all 200 models, and picked only ones consistently ranking above 100 - distribution of ranks was bimodal at the edges and we deemed first mode to end there(see Fig1).

#### 1.3 Coevolution

We used GREMLIN[4] to compute the coevolutionary couplings using DCA (Direct Coupling Analysis), based on PFAM[5] alignment TM7_1 ID: PF00001. We also applied Average Product Correction to the results to alleviate the entropy effects. For the final selection, we standardized the couplings and picked only the pairs above 2 or below -2 standard deviations.

#### 1.4 System preparation

For all systems, we removed all molecules besides the protein. The inserts, fusion proteins, G Protein and antibodies were also removed. Sequences were adjusted to match the main UniProt[6] entries with Modeller[7]: mutations were reversed and short (less than 10 amino acids) gaps were filled. Larger gaps were left as they were. Systems were then embedded in a bilayer containing around 100 POPC molecules divided between the inner and outer leaflet using CHARMM-GUI[8, 9, 10, 11, 12, 13] and all artifical chain ends were capped with amide caps. For all systems, Asp2.50 was protonated, and the protein/membrane assembly placed in a 0.15 M KCl solution. Other acidic amino acids were protonated where environment clearly indicated hydrogen bonding and propka3[14, 15] assigned pKa > 6. All other protonation states were assigned by default for pH 7.

#### 1.5 Simulation setup

All simulations were performed using Gromacs[16, 17, 18, 19, 20, 21, 22], either 2025 or 2026 versions, using the latest release available at the moment of the calculation. We used the CHARMM36m forcefield[23] with TIP3P water model. To equilibrate the systems, we used the default CHARMM-GUI[8, 9, 10, 11, 12, 13] membrane builder protocol of gradually weaker restraints: we ran six simulations of length of 250 ps, 250 ps, 250 ps, 500 ps, 500 ps and 50 ns, respectively. Force constants for position restraints on protein backbone were set to 4000,2000,1000,500,200,50 $\frac{kJ}{mol \cdot nm^2}$, for protein side chain to 2000,1000,500,200,50,0 $\frac{kJ}{mol \cdot nm^2}$, and for lipid phosphorus atoms to 1000,400,400,200,40,0 $\frac{kJ}{mol \cdot nm^2}$, respectively, for 6 consecutive simulations. Force constants for lipid dihedral restraints were set to 1000,400,200,200,100,0 $\frac{kJ}{mol \cdot rad^2}$, respectively, for C1-C3-C2-O21 and C28-C29-C210-C211 dihedrals. Afterwards, equilibrium runs have been run for 100 ns, and trajectories were deposited every 50 ps. These runs were then used for RMSD comparisons and to train an autoencoder. All simulations were carried using a timestep of 2 fs, a temperature set to 310K and a pressure set to 1 bar, the v-rescale thermostat with tau 1.0 and the C-rescale barostat[24] with tau 5.0, semiisotropic coupling and nstpcouple set to 10. Hydrogen bonds were constrained using the linear constraint solver (LINCS)[25], and long-range electrostatics were accounted for using the particle mesh Ewald (PME) method beyond the 12 Åelectrostatic cutoff. A neighborlist cutoff was used for vdw interactions with rvdw equal 12 Åand a switching function starting at 10 Å.

#### 1.6 Biasing protocol

For transition simulations, final frames of equilibration in either state were chosen at random as a starting point. The simulations were then biased along the general collective variables selected as described above using Accelerated Weight Histogram (AWH)[26, 27, 28] as implemented in GROMACS[16, 17, 18, 19, 20, 21, 22]. We used Boltzmann as a target, with beta scaling set to 0.01, awh-nstsample set to 500, equilibrate histogram set on, sampling dimensions set from -2.5 to -0.4 for the first collective variable and -0.1 to 1.0 for the second collective variable, force constants 10000 and 5000, respectively, and diffusion equal to 0.000005. There were a few exceptions to these parameters: the second dimension was extended to -0.2 for the canabinoid receptor, -0.4 for the chemokine receptor, and -0.3 for opsins. For prostanoid, S1P, somatostatin, neuropeptide Y and neurotensin receptors, beta scaling was set to 0.05 and the diffusion set to 0.0000085 and 0.0000012 for CV1 and CV2, respectively. For all systems and all runs, each individual distance that was biased as a part of either CV was restrained with flat-bottom (one-sided harmonic) potential centered at the highest value + 10% of that value of that distance present in the training set, i.ex. highest value of distance present in the PDB structures. All simulations were run using a single walker. Simulations were run until multiple coverings occurred, and data only up until first covering was used for analyses unless indicated otherwise. Trajectories were deposited every 100 ps.

#### 1.7 RMSD

To validate our transitions, we have run 100 ns equilibrium, unbiased MD simulations of each receptor starting from active and inactive structures. As these structures differed in the resolved parts in all cases, we first aligned their sequences, and then picked only helical parts for RMSD computations. As helicity could also differ between the structures, the amino acid selection in each receptor has been manually adjusted to match. We provide final selection of residues as PDB files for each receptor. For each frame from our transition trajectories, we computed the C$\alpha$ RMSD of aforementioned selections to each frame in active and inactive equilibrium, respectively. For each frame from transitions, we then report a mean of RMSD to all equilibrium frames in the respective states.

#### 1.8 Microswitches

To have a common frame of reference for microswitches, we sought to describe them with invariant measures. Therefore, we defined CWxP as $\chi_2$ of TRP/TYR/PHE; NPxxY as distance between C$\alpha$ of amino acids positions corresponding to N and Y; TM5 bulge as distance between C$\alpha$ of amino acids at alignment positions corresponding corresponding to GLY315 and SER207 in the $\beta$2-adrenergic receptor; the connector as the dihedral angle between C$\beta$, C$\alpha$, C$\alpha$, C$\beta$ of amino acids at alignment positions corresponding to PHE282 and LEU121 in the $\beta$2-adrenergic receptor; DRY as the distance between C$\alpha$ and C$\zeta$ of arginine, and TM3-TM6 distance as the distance between the C$\alpha$ of amino acids at alignment positions corresponding to ARG131 and and LYS270 in the $\beta$2-adrenergic receptor. We provide transition trajectories for the readers to assess structural changes themselves as a part of this manuscript.

#### 1.9 Secondary structure

To show that secondary structure has not changed in a significant manner during biasing simulations, we have computed the percentage of coil, helix and sheet in the transition trajectories using the MDtraj[29] DSSP function. Dashed lines indicate transition times.

#### 1.10 Autoencoder

The autoencoder[30] was composed of layers of sizes [450, 300, 150, 50, 25, 10, 2, 10, 25, 50, 150, 300, 450], with LeakyReLU activation functions with slope set to 0.2. It was trained using AdamW with weight decay equal to 0.001 and L1 function as loss, with training batch size of 50000 and validation batch size 5000. For each batch, we sampled equal amounts of each of the 20 receptors. We also used Mutltistep Learning Rate scheduler that decreased the learning rate 10 times every 2500 steps. Training was stopped early at epoch 7215, with early stopping parametrized with patience 2500 and delta equal 0.0001. Only equilibrium simulations were used for training, and GPR183 was excluded. For feature importance analysis, we used layerwise relevance propagation[31] as SHAP[3] was prohibitively expensive for the size of the model and the dataset. We ranked all the feature importances for each system, and picked only the ones in top 100 for further consideration.

#### 1.11 Free energy profiles

For $\beta$2-adrenergic receptor free energy surfaces(FES), ligands have been copy-pasted from their experimental structures 6PS0(carazolol)[32], 6PS6(timolol)[32], 6MXT(salmeterol)[33], 7BZ2(formoterol)[34], 3NYA(alprenolol)[35] to 4LDO and equilibrated in the same way as described in System preparation.[36] We used the adrenaline bound to 4LDO for simulations bound with adrenaline.[37] We used two first principal components of the $\beta$2-adrenergic receptor transition, only considering the distances present in the common CVs. Prior to PCA, all distances were transformed in the following way (this transformation was also used for the final CV):

$$e^{-\frac{x-min}{max-min}}$$

instead of previously used exponential differences. Depending on the system, 6 to 12 copies of biasing with AWH[26, 27, 28] have been run, with awh-nstsample set to 100(with exception of carazolol, for which it was set to 1000), target set to constant, equilibrate histogram set on, CVs ranging from -1.84 to 0.12 and -1.69 to -0.52 for first and second PC, respectively. Force constants were set to 7500 and diffusion was set to 0.00005 for both dimensions. Cover diameter was left as default 0.15. Means of last frames from all copies were reported as final free energy surfaces. Every copy was run with 4 walkers, and reported simulation times are per walker. If a copy did not leave the initial stage in the two rounds of simulations (e.g. 6h job on the cluster) it was discarded without checking the result and the copy was re-set up. Convergence was assessed by the distances between the final FES and estimates of FES by each copy in time. Outlier copies, as indicated by IQR analysis, have been excluded from analysis and final FES computation. These copies are indicated on the figures by the gray coloring. Errors for each FES were computed with unbiased jackknife standard deviation estimator:

$$\sqrt{\frac{N-1}{N} \cdot \sum_i (x_i - \bar{x})^2}$$

Table 1: Frequencies in % of amino acids in the multiple sequence alignment of selected conserved positions. BW numbering was matched to internal MSA numbering based on the $\beta$2-adrenergic receptor. Frequencies below 1% and gaps have been omitted.

| 6x48 | 5x58 | 7x49 | 3x39 | 7x45 |
|---|---|---|---|---|
| W 57.1 | Y 67.7 | N 63.7 | S 68.6 | N 51.7 |
| F 9.9 | N 9.4 | D 10.2 | T 6.4 | S 13.1 |
| Y 4.4 | S 3.1 | | G 4.8 | H 4.6 |
| L 1.8 | T 1.6 | | A 3.3 | A 1.5 |
| N 1.6 | H 1.4 | | | K 1.4 |
| S 1.6 | F 1.1 | | | Q 1.0 |
| Q 1.6 | | | | T 1.0 |
| T 1.4 | | | | |
| M 1.2 | | | | |
| V 1.1 | | | | |

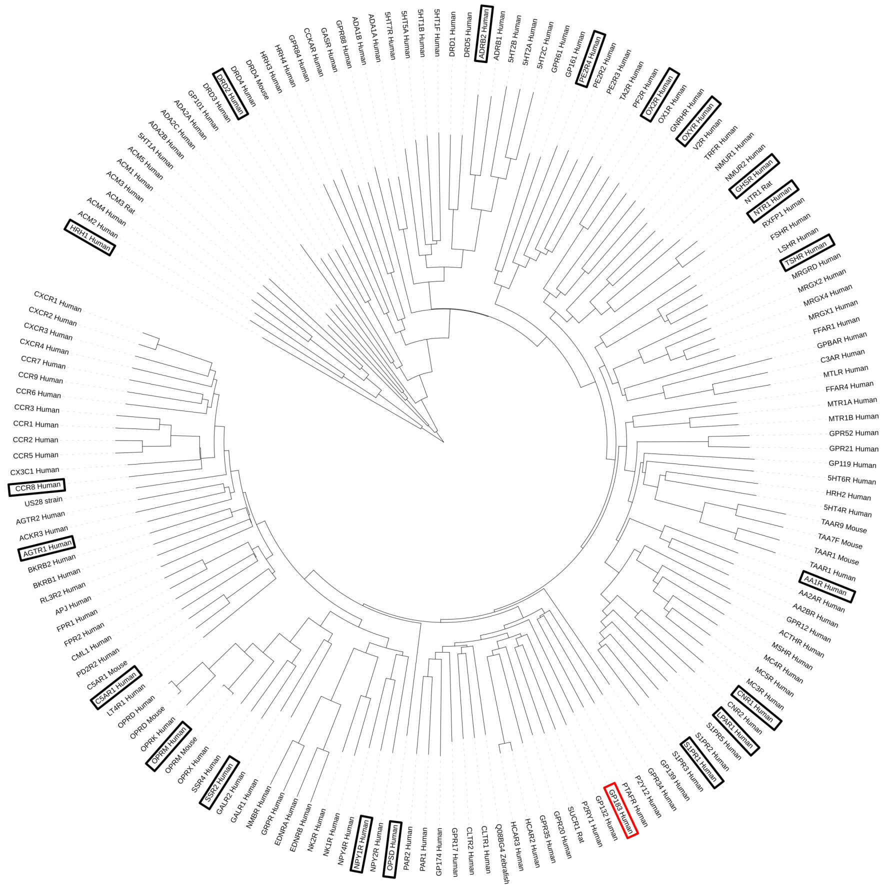

Figure 1: Phylogenetic tree of all receptors from our training set (GPCRdb[1] from 11.2022). Receptors selected as representatives for MD simulations are highlighted with a black frame, and GPR183 is highlighted with a red frame. The tree was computed using ParsimonyTreeConstructor from Biopython[75] with all default options, and visualized with iToL.[76]

Table 2: PDB codes and simulation times for all systems we have simulated. Bold font indicates state which we have started transition from. Some of these structures have been obtained and added to GPCRdb[1] after 11.2022 - these were never used in the process of deriving CVs. Simulation time marks first covering in the AWH algorithm, trajectories were not analyzed beyond it.

| *Receptor family*<br>Time to transition | *Gene name*<br>Simulation time | *Inactive pdb* | *Active pdb* |
|---|---|---|---|
| chemokine<br>1994 ns | CCR8 | **8U1U** [38] | 8TLM[38] 838 ns |
| cannabinoid<br>1899 ns | CNR1 | 8GHV[39] | **5U09**[40] 348 ns |
| prostanoid<br>480 ns | PE2R4<br>1282 ns | 8GDB[41] | **5YWY**[42] |
| opioid<br>458 ns | OPRM<br>1259 ns | 8F7Q[43] | **7UL4**[44] |
| complement peptide<br>302 ns | C5AR1<br>1607 ns | **7Y65**[45] | 8HK5[46] |
| somatostatin<br>56 ns | SSR2<br>1115 ns | **7Y27**[47] | 7XNA[48] |
| neuropeptide Y<br>338 ns | NPY1R<br>1315 ns | 7X9A[49] | **5ZBQ**[50] |
| ghrelin<br>354 ns | GHSR<br>883 ns | **7W2Z**[51] | 6KO5[52] |
| S1P<br>60 ns | S1PR1<br>958 ns | **7TD3**[53] | 3V2W[54] |
| LPA<br>395 ns | LPAR1<br>974 ns | **7TD0**[53] | 4Z36[55] |
| glycoprotein hormone<br>590 ns | TSHR<br>998 ns | 7T9I[56] | **7T9M**[56] |
| VaO<br>595 ns | OXYR<br>1899 ns | **7RYC**[57] | 6TPK[58] |
| adenosine<br>298 ns | AA1R<br>1039 ns | **7LD4**[59] | 5UEN[60] |
| orexin<br>119 ns | OX2R<br>1017 ns | **7L1U**[61] | 5WQC [62] |
| dopamine<br>490 ns | DRD2<br>1254 ns | 7JVR[63] | **6CM4**[64] |
| angiotensin<br>201 ns | AGTR1<br>939 ns | 7F6G[65] | **4ZUD**[66] |
| histamine<br>265 ns | HRH1<br>1696 ns | **7DFL**[67] | 3RZE[68] |
| opsins<br>596 ns | OPSD<br>1770 ns | 6OY9[69] | **8A6C**[70] |
| neurotensin<br>568 ns | NTR1<br>1846 ns | **6OS9**[71] | 4BUO[72] |
| adrb2<br>18 ns | ADRB2<br>1199 ns | **4LDO**[37] | 2RH1[73] |
| GPR183<br>593 ns | GPR183<br>953 ns | **7TUZ**[74] | 7TUY[74] |

Table 3: Protonatonated residues besides Asp 2.50 . For all these environment clearly indicated hydrogen bonding and propka3[14, 15] assigned pKa > 6

| **system** | textbfadditional protonated residues |
|---|---|
| prostanoid | ASP325<br>GLU 116 |
| somatostatin | GLU90 |
| opsins | GLU122<br>GLU181 |
| glycoprotein hormone | ASP633<br>GLU506 |
| cannabinoid | GLU133 |
| VaO | ASP153 |
| dopamine | GLU62 |
| adrb2 | GLU122 |
| GPR183 | ASP304 |
| S1P active | GLU62 |

#### 1.12 chemokine

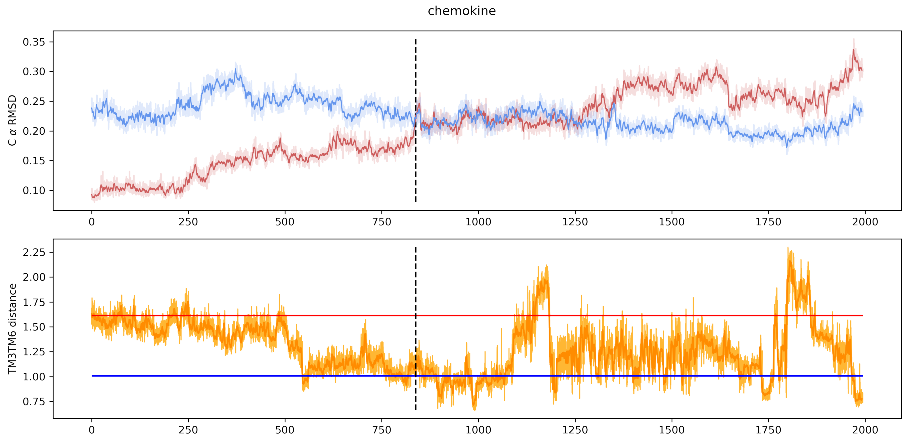


Figure 2: RMSD traces to active and inactive equilibrium ensembles (top) and TM3TM6 distance (bottom) for transition trajectory for chemokine system. On both plots, running mean of size 25 (2.5 ns) shown in darker shades, and values for particular frames shown in lighter shades. Mean values of TM3TM6 distances from active (red) and inactive (blue) equilibrium simulations are shown as horizontal lines.

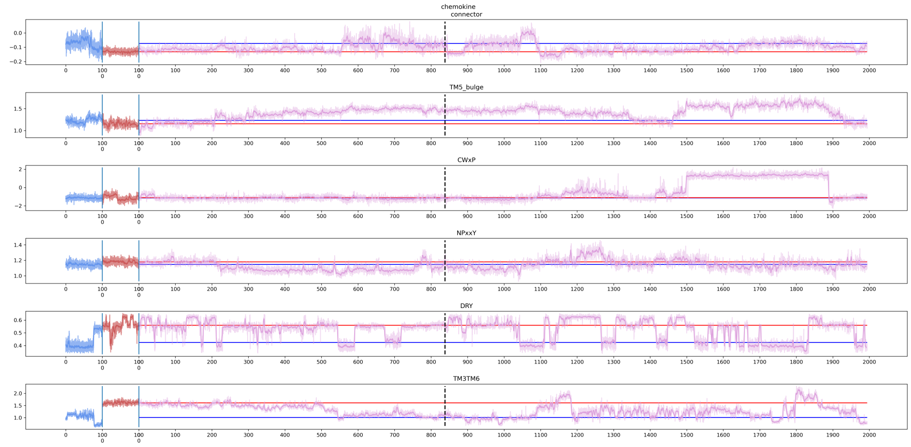


Figure 3: Selected microswitches changes in time for chemokine system. On all plots, running mean of size 25 (2.5 ns, 1.25 ns for equilibria) shown in darker shades, and values for particular frames shown in lighter shades. Values for active (red) and inactive (blue) equilibrium simulations are shown at the beginning of each microswitch plot. Values for transition trajectory (purple) follows. Mean values from equilibrium simulations are shown as red (active) and blue (inactive) horizontal lines.

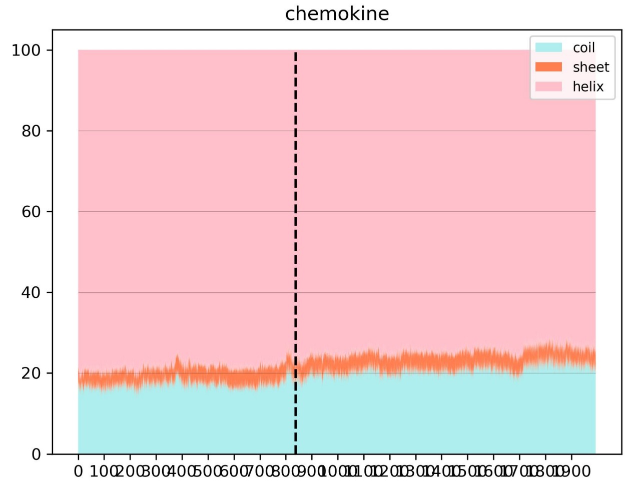


Figure 4: Secondary structure elements percentage for transition trajectory for chemokine system.

#### 1.13 cannabinoid

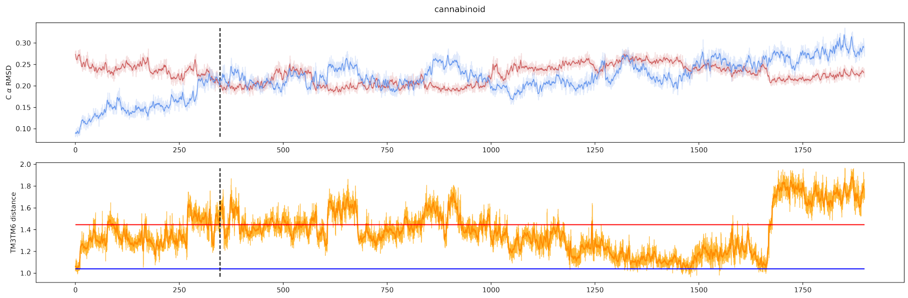


Figure 5: RMSD traces to active and inactive equilibrium ensembles (top) and TM3TM6 distance (bottom) for transition trajectory for cannabinoid system. On both plots, running mean of size 25 (2.5 ns) shown in darker shades, and values for particular frames shown in lighter shades. Mean values of TM3TM6 distances from active (red) and inactive (blue) equilibrium simulations are shown as horizontal lines.

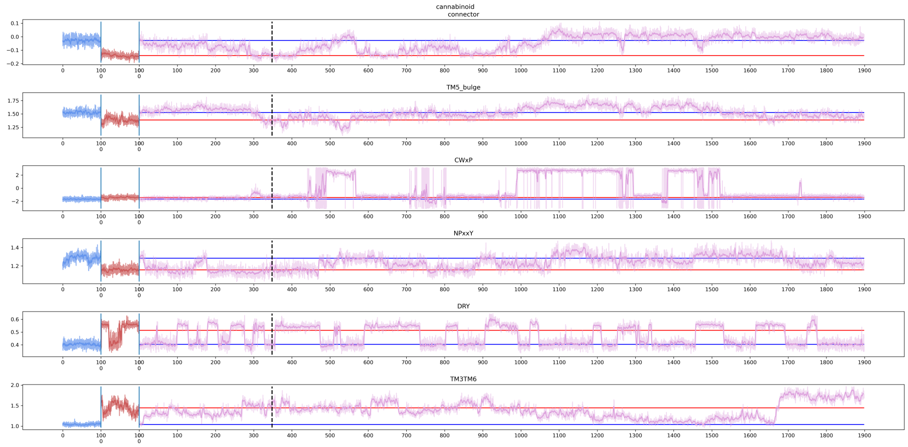


Figure 6: Selected microswitches changes in time for cannabinoid system. On all plots, running mean of size 25 (2.5 ns, 1.25 ns for equilibria) shown in darker shades, and values for particular frames shown in lighter shades. Values for active (red) and inactive (blue) equilibrium simulations are shown at the beginning of each microswitch plot. Values for transition trajectory (purple) follows. Mean values from equilibrium simulations are shown as red (active) and blue (inactive) horizontal lines.

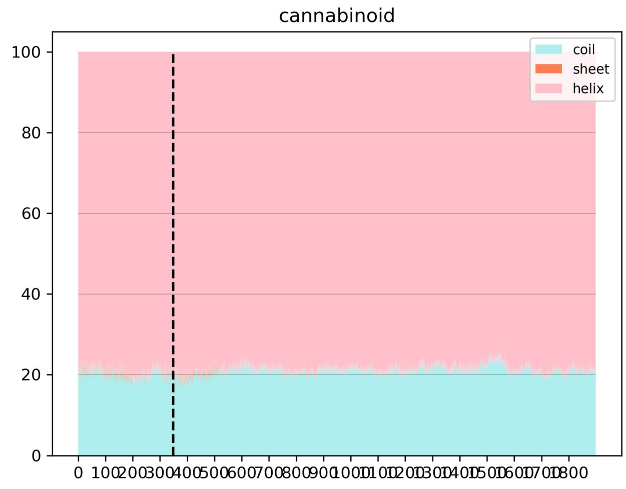


Figure 7: Secondary structure elements percentage for transition trajectory for cannabinoid system.

#### 1.14 prostanoid

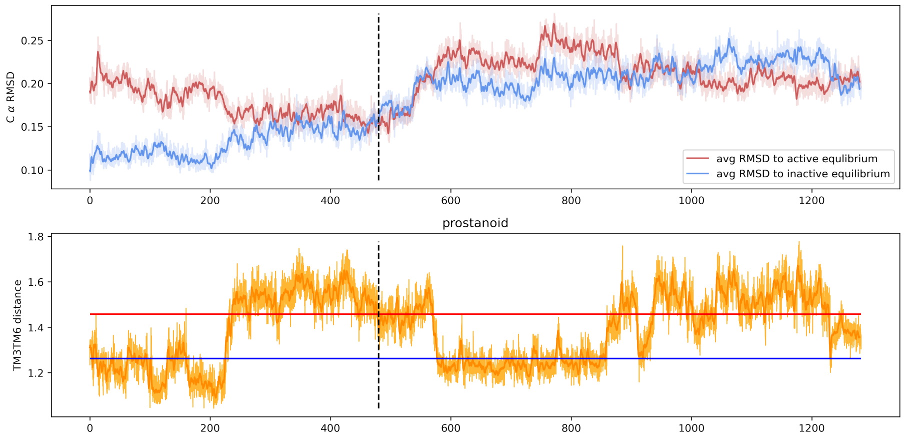


Figure 8: RMSD traces to active and inactive equilibrium ensembles (top) and TM3TM6 distance (bottom) for transition trajectory for prostanoid system. On both plots, running mean of size 25 (2.5 ns) shown in darker shades, and values for particular frames shown in lighter shades. Mean values of TM3TM6 distances from active (red) and inactive (blue) equilibrium simulations are shown as horizontal lines.

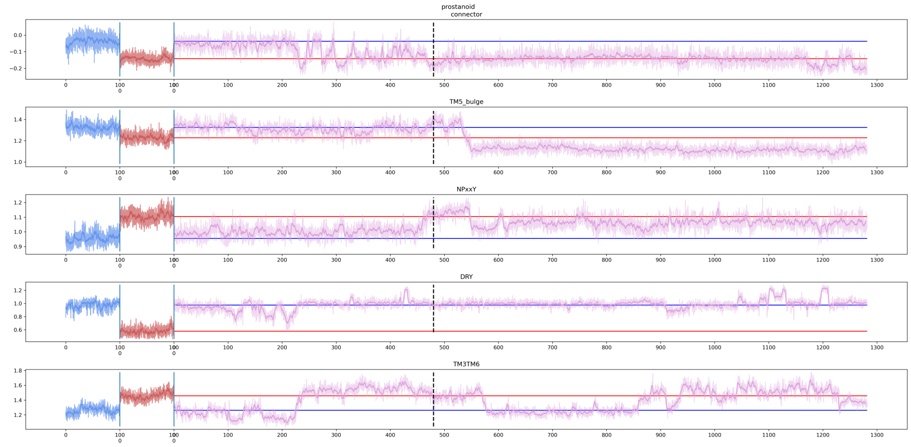


Figure 9: Selected microswitches changes in time for prostanoid system. On all plots, running mean of size 25 (2.5 ns, 1.25 ns for equilibria) shown in darker shades, and values for particular frames shown in lighter shades. Values for active (red) and inactive (blue) equilibrium simulations are shown at the beginning of each microswitch plot. Values for transition trajectory (purple) follows. Mean values from equilibrium simulations are shown as red (active) and blue (inactive) horizontal lines.

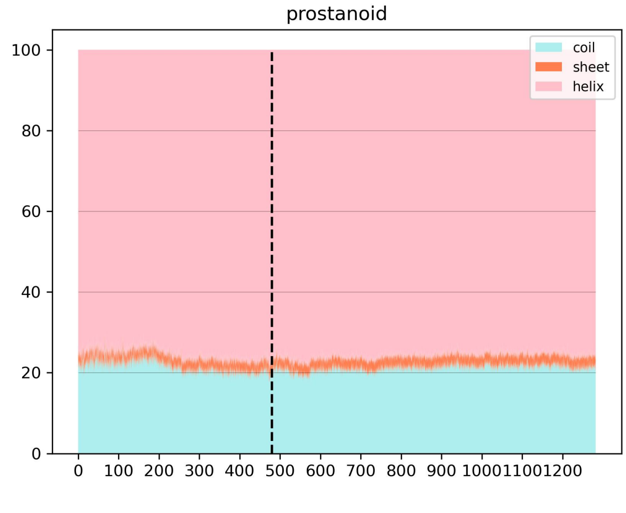


Figure 10: Secondary structure elements percentage for transition trajectory for prostanoid system.

#### 1.15 opioid

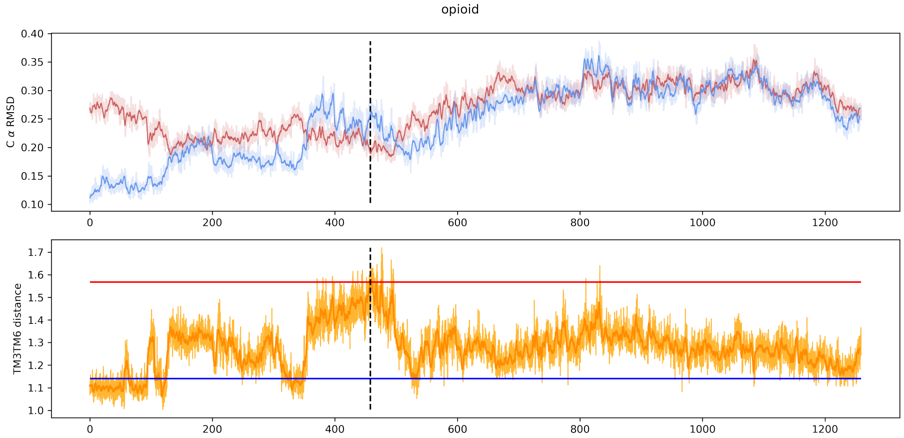


Figure 11: RMSD traces to active and inactive equilibrium ensembles (top) and TM3TM6 distance (bottom) for transition trajectory for opioid system. On both plots, running mean of size 25 (2.5 ns) shown in darker shades, and values for particular frames shown in lighter shades. Mean values of TM3TM6 distances from active (red) and inactive (blue) equilibrium simulations are shown as horizontal lines.

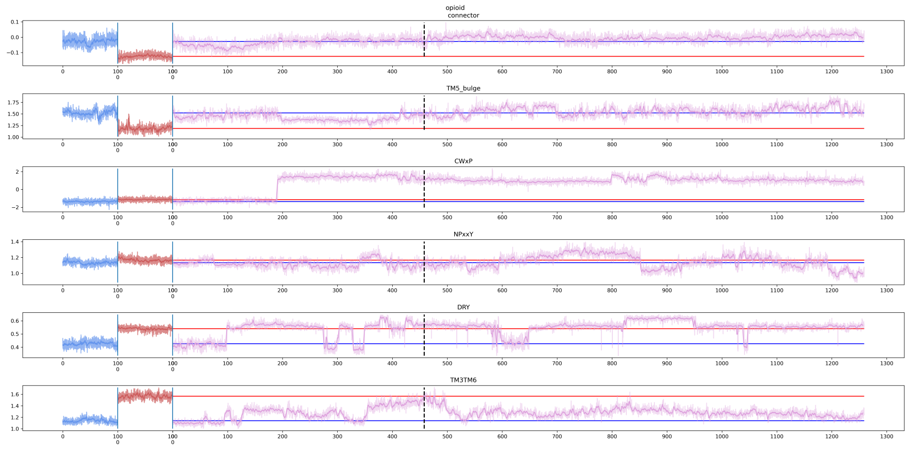


Figure 12: Selected microswitches changes in time for opioid system. On all plots, running mean of size 25 (2.5 ns, 1.25 ns for equilibria) shown in darker shades, and values for particular frames shown in lighter shades. Values for active (red) and inactive (blue) equilibrium simulations are shown at the beginning of each microswitch plot. Values for transition trajectory (purple) follows. Mean values from equilibrium simulations are shown as red (active) and blue (inactive) horizontal lines.

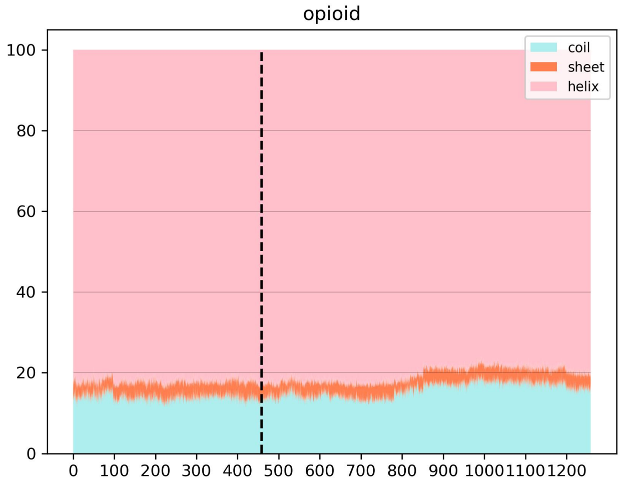


Figure 13: Secondary structure elements percentage for transition trajectory for opioid system.

#### 1.16 complement peptide

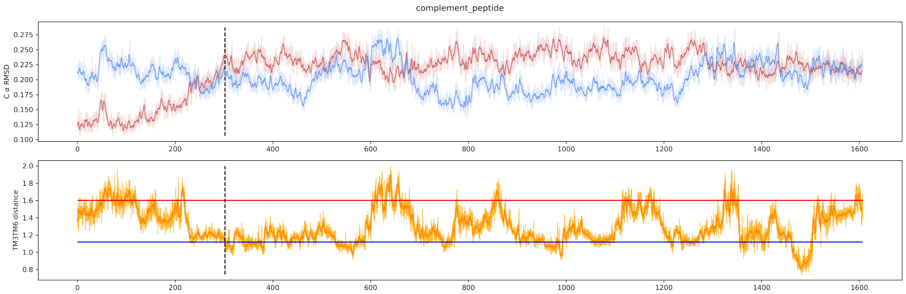


Figure 14: RMSD traces to active and inactive equilibrium ensembles (top) and TM3TM6 distance (bottom) for transition trajectory for complement peptide system. On both plots, running mean of size 25 (2.5 ns) shown in darker shades, and values for particular frames shown in lighter shades. Mean values of TM3TM6 distances from active (red) and inactive (blue) equilibrium simulations are shown as horizontal lines.

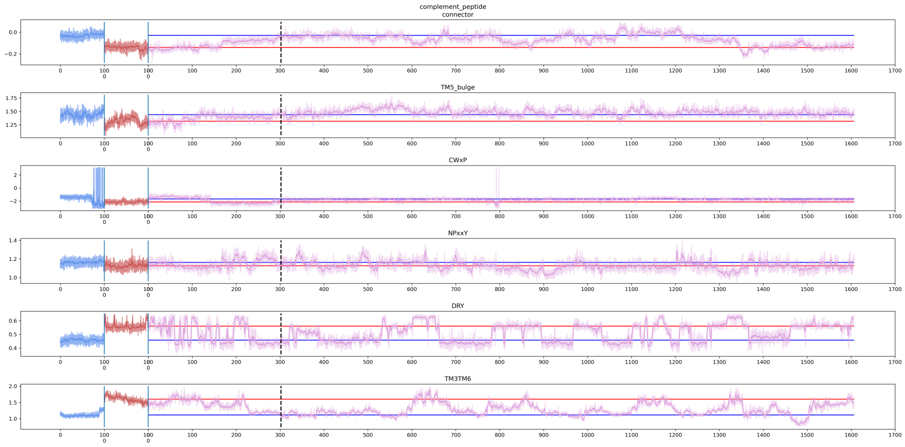


Figure 15: Selected microswitches changes in time for complement peptide system. On all plots, running mean of size 25 (2.5 ns, 1.25 ns for equilibria) shown in darker shades, and values for particular frames shown in lighter shades. Values for active (red) and inactive (blue) equilibrium simulations are shown at the beginning of each microswitch plot. Values for transition trajectory (purple) follows. Mean values from equilibrium simulations are shown as red (active) and blue (inactive) horizontal lines.

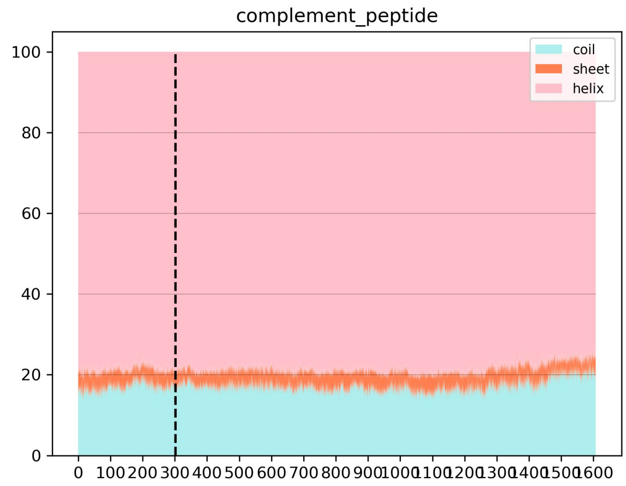


Figure 16: Secondary structure elements percentage for transition trajectory for complement peptide system.

#### 1.17 somatostatin

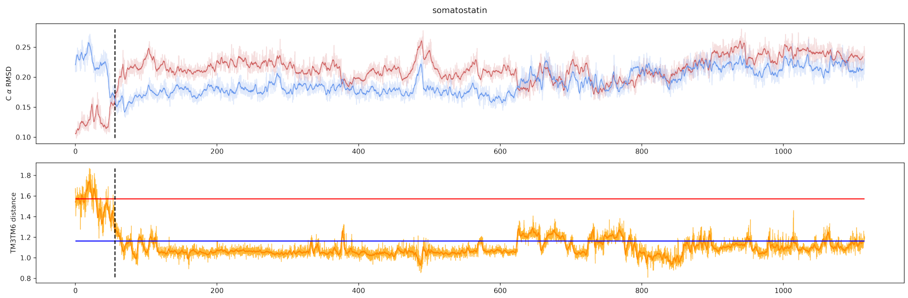


Figure 17: RMSD traces to active and inactive equilibrium ensembles (top) and TM3TM6 distance (bottom) for transition trajectory for somatostatin system. On both plots, running mean of size 25 (2.5 ns) shown in darker shades, and values for particular frames shown in lighter shades. Mean values of TM3TM6 distances from active (red) and inactive (blue) equilibrium simulations are shown as horizontal lines.

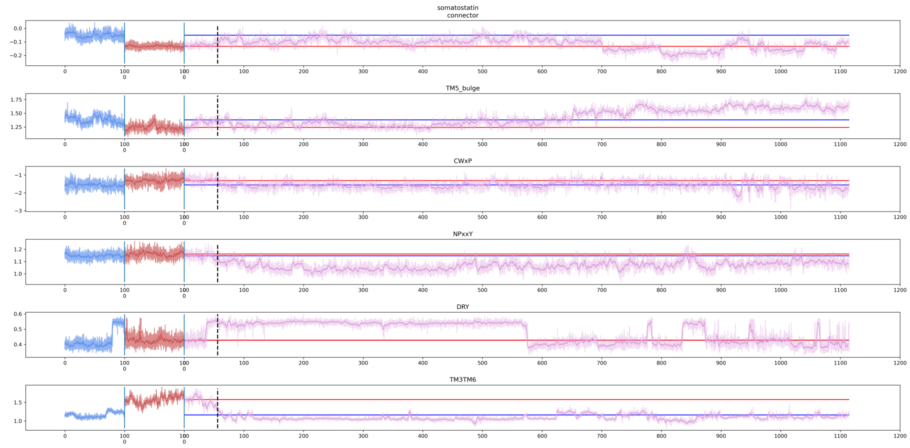


Figure 18: Selected microswitches changes in time for somatostatin system. On all plots, running mean of size 25 (2.5 ns, 1.25 ns for equilibria) shown in darker shades, and values for particular frames shown in lighter shades. Values for active (red) and inactive (blue) equilibrium simulations are shown at the beginning of each microswitch plot. Values for transition trajectory (purple) follows. Mean values from equilibrium simulations are shown as red (active) and blue (inactive) horizontal lines.

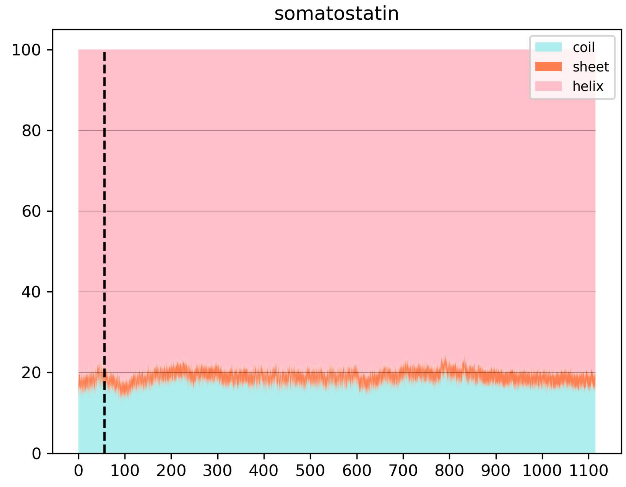


Figure 19: Secondary structure elements percentage for transition trajectory for somatostatin system.

#### 1.18 neuropeptide Y

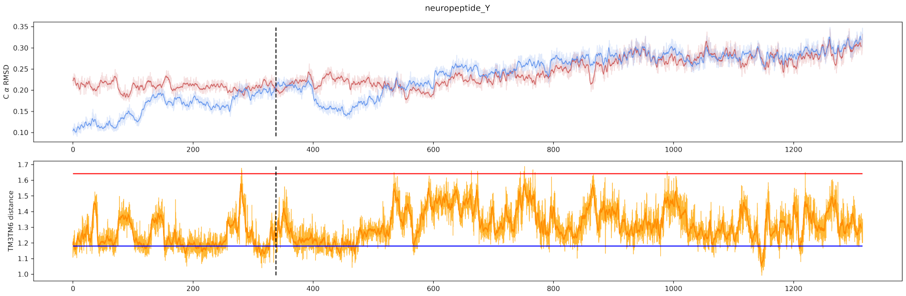


Figure 20: RMSD traces to active and inactive equilibrium ensembles (top) and TM3TM6 distance (bottom) for transition trajectory for neuropeptide Y system. On both plots, running mean of size 25 (2.5 ns) shown in darker shades, and values for particular frames shown in lighter shades. Mean values of TM3TM6 distances from active (red) and inactive (blue) equilibrium simulations are shown as horizontal lines.

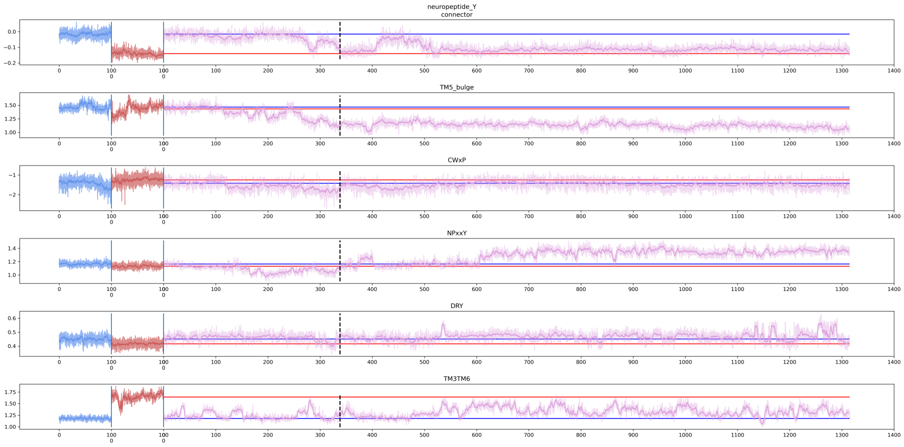


Figure 21: Selected microswitches changes in time for neuropeptide Y system. On all plots, running mean of size 25 (2.5 ns, 1.25 ns for equilibria) shown in darker shades, and values for particular frames shown in lighter shades. Values for active (red) and inactive (blue) equilibrium simulations are shown at the beginning of each microswitch plot. Values for transition trajectory (purple) follows. Mean values from equilibrium simulations are shown as red (active) and blue (inactive) horizontal lines.

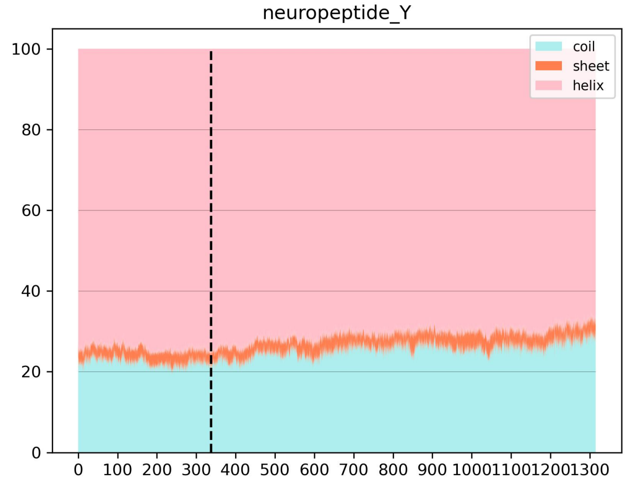


Figure 22: Secondary structure elements percentage for transition trajectory for neuropeptide Y system.

#### 1.19 ghrelin

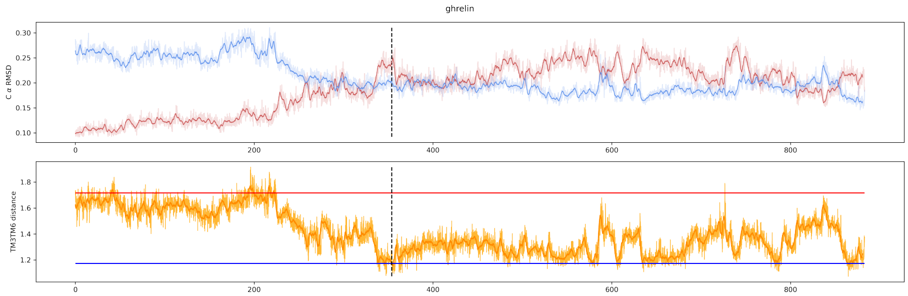


Figure 23: RMSD traces to active and inactive equilibrium ensembles (top) and TM3TM6 distance (bottom) for transition trajectory for ghrelin system. On both plots, running mean of size 25 (2.5 ns) shown in darker shades, and values for particular frames shown in lighter shades. Mean values of TM3TM6 distances from active (red) and inactive (blue) equilibrium simulations are shown as horizontal lines.

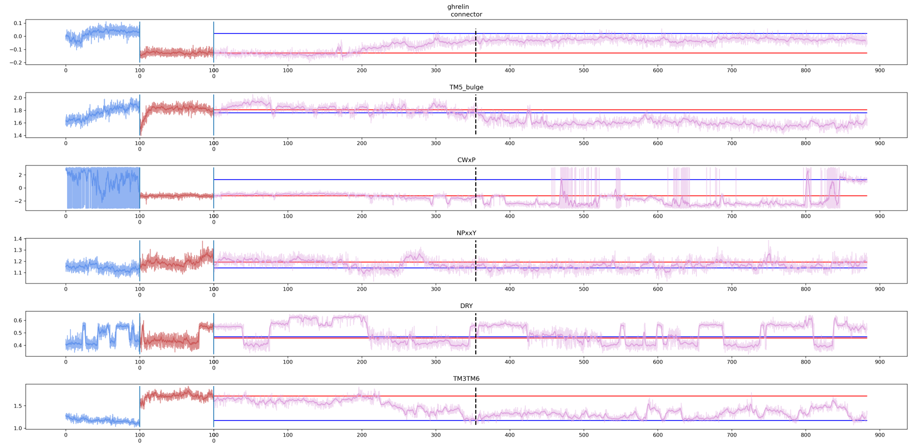


Figure 24: Selected microswitches changes in time for ghrelin system. On all plots, running mean of size 25 (2.5 ns, 1.25 ns for equilibria) shown in darker shades, and values for particular frames shown in lighter shades. Values for active (red) and inactive (blue) equilibrium simulations are shown at the beginning of each microswitch plot. Values for transition trajectory (purple) follows. Mean values from equilibrium simulations are shown as red (active) and blue (inactive) horizontal lines.

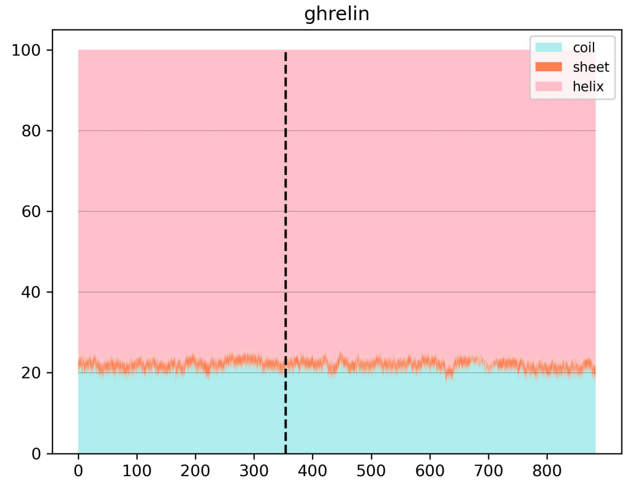


Figure 25: Secondary structure elements percentage for transition trajectory for ghrelin system.

## 1.20 S1P

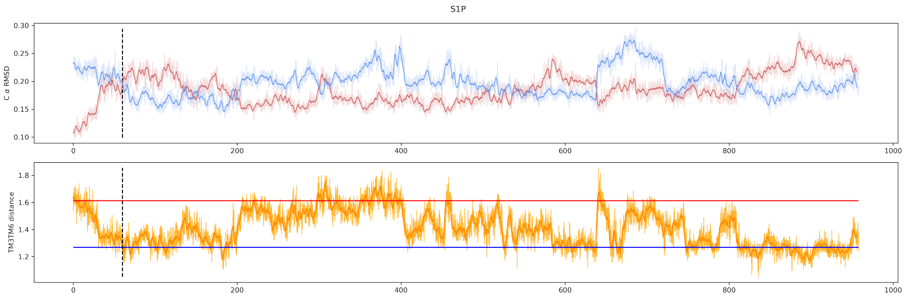


Figure 26: RMSD traces to active and inactive equilibrium ensembles (top) and TM3TM6 distance (bottom) for transition trajectory for S1P system. On both plots, running mean of size 25 (2.5 ns) shown in darker shades, and values for particular frames shown in lighter shades. Mean values of TM3TM6 distances from active (red) and inactive (blue) equilibrium simulations are shown as horizontal lines.

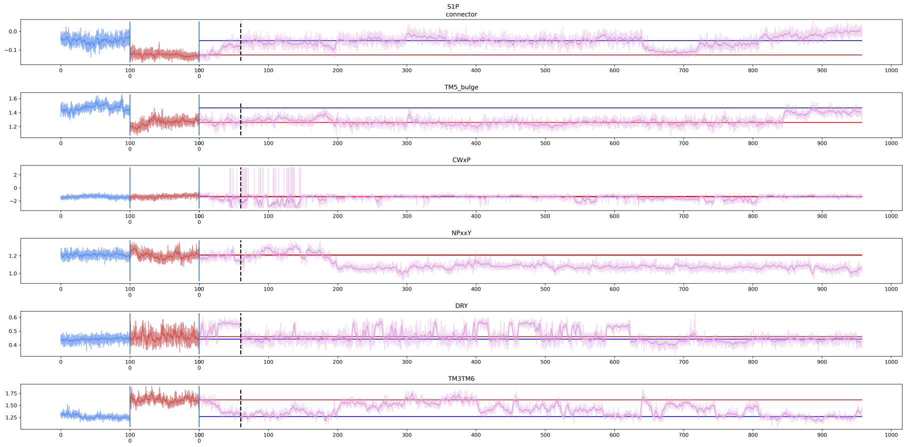


Figure 27: Selected microswitches changes in time for S1P system. On all plots, running mean of size 25 (2.5 ns, 1.25 ns for equilibria) shown in darker shades, and values for particular frames shown in lighter shades. Values for active (red) and inactive (blue) equilibrium simulations are shown at the beginning of each microswitch plot. Values for transition trajectory (purple) follows. Mean values from equilibrium simulations are shown as red (active) and blue (inactive) horizontal lines.

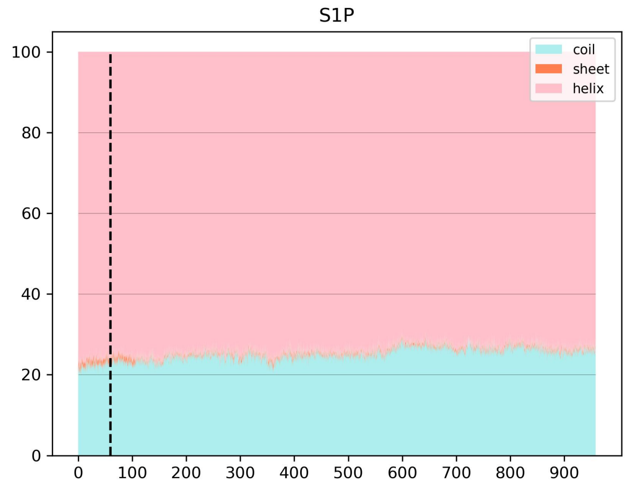


Figure 28: Secondary structure elements percentage for transition trajectory for S1P system.

#### 1.21 LPA

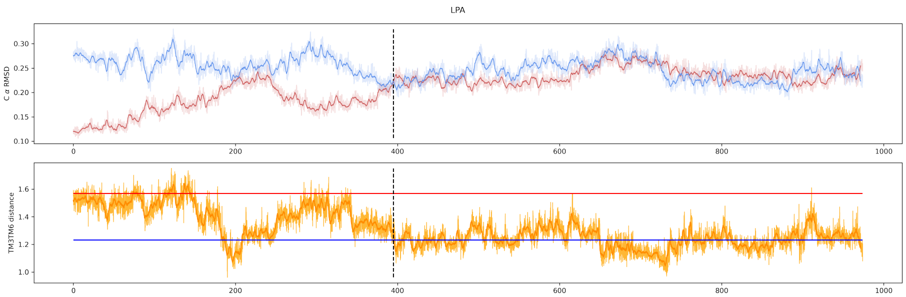


Figure 29: RMSD traces to active and inactive equilibrium ensembles (top) and TM3TM6 distance (bottom) for transition trajectory for LPA system. On both plots, running mean of size 25 (2.5 ns) shown in darker shades, and values for particular frames shown in lighter shades. Mean values of TM3TM6 distances from active (red) and inactive (blue) equilibrium simulations are shown as horizontal lines.

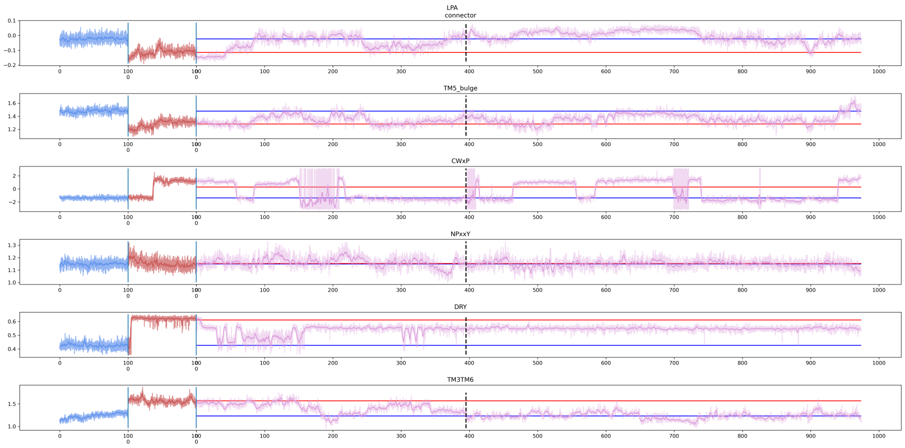


Figure 30: Selected microswitches changes in time for LPA system. On all plots, running mean of size 25 (2.5 ns, 1.25 ns for equilibria) shown in darker shades, and values for particular frames shown in lighter shades. Values for active (red) and inactive (blue) equilibrium simulations are shown at the beginning of each microswitch plot. Values for transition trajectory (purple) follows. Mean values from equilibrium simulations are shown as red (active) and blue (inactive) horizontal lines.

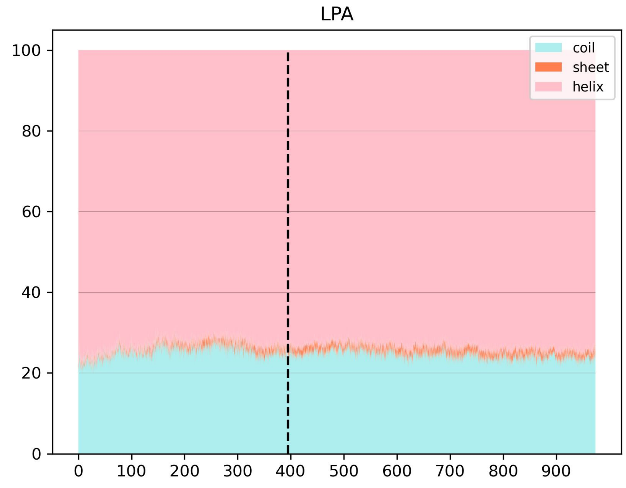


Figure 31: Secondary structure elements percentage for transition trajectory for LPA system.

#### 1.22 glycoprotein hormone

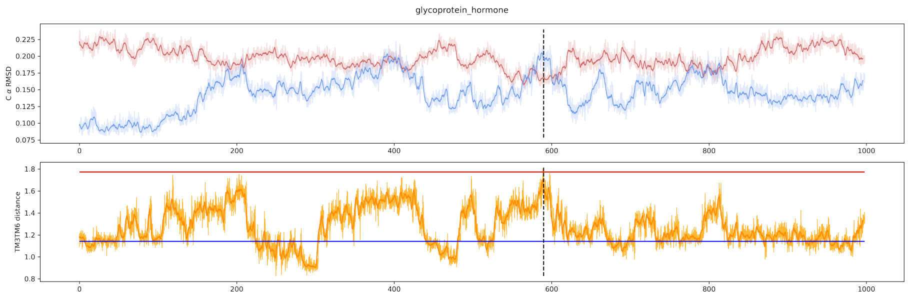


Figure 32: RMSD traces to active and inactive equilibrium ensembles (top) and TM3TM6 distance (bottom) for transition trajectory for glycoprotein hormone system. On both plots, running mean of size 25 (2.5 ns) shown in darker shades, and values for particular frames shown in lighter shades. Mean values of TM3TM6 distances from active (red) and inactive (blue) equilibrium simulations are shown as horizontal lines.

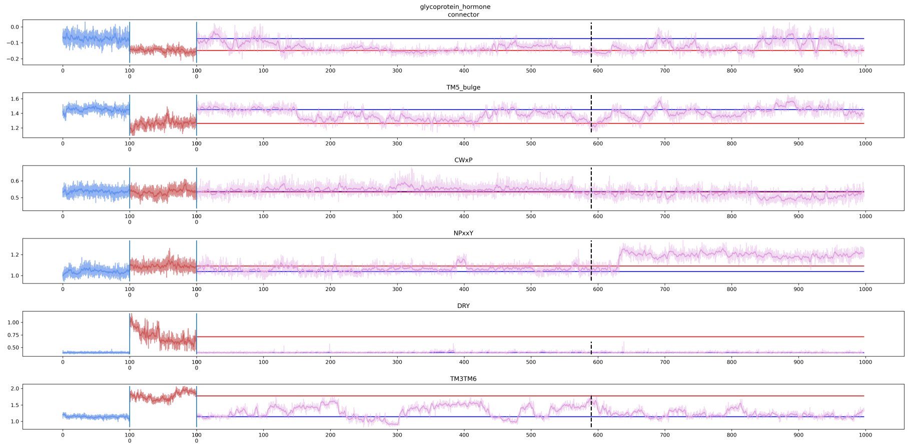


Figure 33: Selected microswitches changes in time for glycoprotein hormone system. On all plots, running mean of size 25 (2.5 ns, 1.25 ns for equilibria) shown in darker shades, and values for particular frames shown in lighter shades. Values for active (red) and inactive (blue) equilibrium simulations are shown at the beginning of each microswitch plot. Values for transition trajectory (purple) follows. Mean values from equilibrium simulations are shown as red (active) and blue (inactive) horizontal lines.

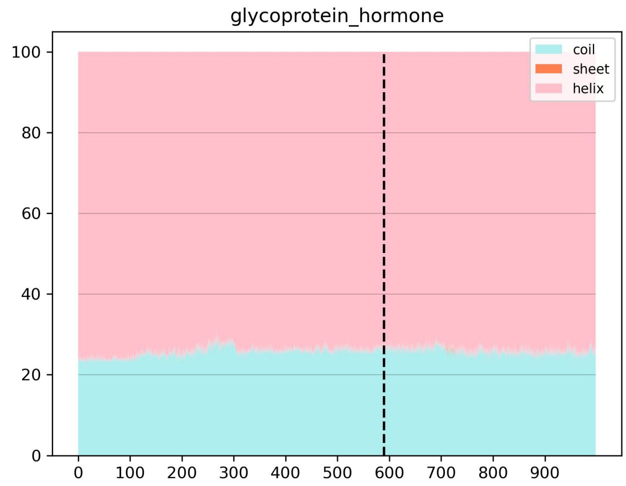


Figure 34: Secondary structure elements percentage for transition trajectory for glycoprotein hormone system.

#### 1.23 VaO

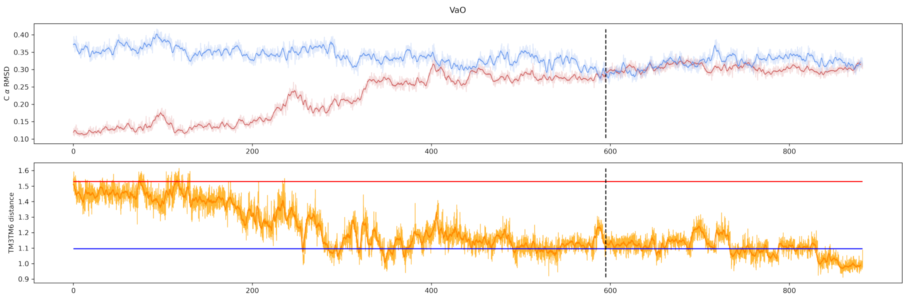


Figure 35: RMSD traces to active and inactive equilibrium ensembles (top) and TM3TM6 distance (bottom) for transition trajectory for VaO system. On both plots, running mean of size 25 (2.5 ns) shown in darker shades, and values for particular frames shown in lighter shades. Mean values of TM3TM6 distances from active (red) and inactive (blue) equilibrium simulations are shown as horizontal lines.

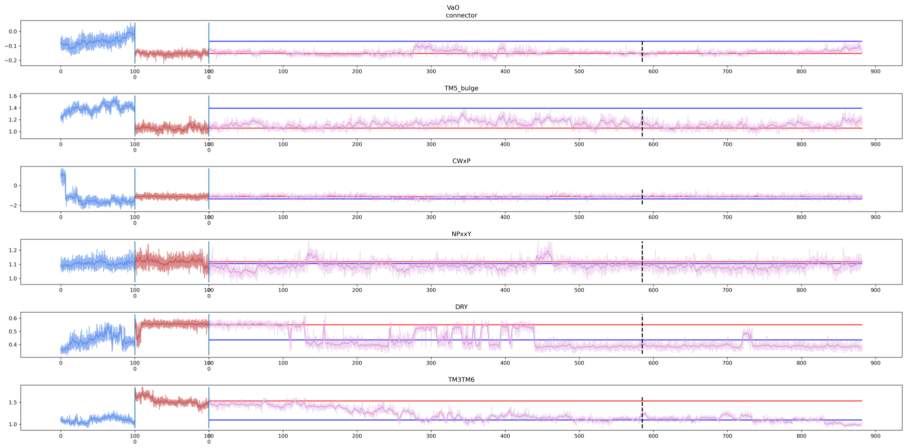


Figure 36: Selected microswitches changes in time for VaO system. On all plots, running mean of size 25 (2.5 ns, 1.25 ns for equilibria) shown in darker shades, and values for particular frames shown in lighter shades. Values for active (red) and inactive (blue) equilibrium simulations are shown at the beginning of each microswitch plot. Values for transition trajectory (purple) follows. Mean values from equilibrium simulations are shown as red (active) and blue (inactive) horizontal lines.

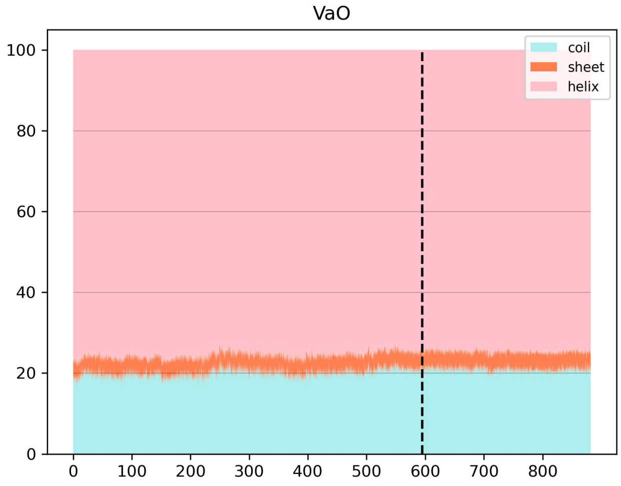


Figure 37: Secondary structure elements percentage for transition trajectory for VaO system.

#### 1.24 adenosine

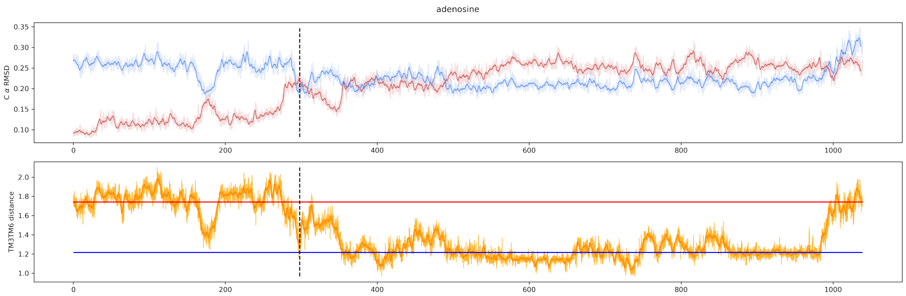


Figure 38: RMSD traces to active and inactive equilibrium ensembles (top) and TM3TM6 distance (bottom) for transition trajectory for adenosine system. On both plots, running mean of size 25 (2.5 ns) shown in darker shades, and values for particular frames shown in lighter shades. Mean values of TM3TM6 distances from active (red) and inactive (blue) equilibrium simulations are shown as horizontal lines.

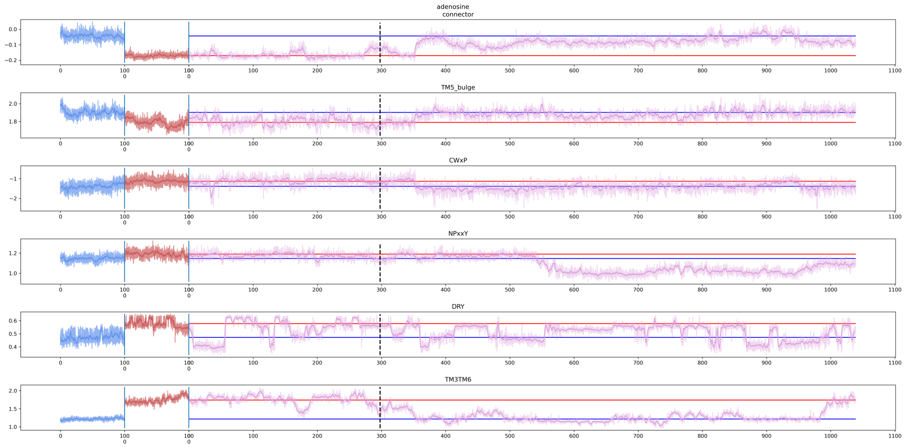


Figure 39: Selected microswitches changes in time for adenosine system. On all plots, running mean of size 25 (2.5 ns, 1.25 ns for equilibria) shown in darker shades, and values for particular frames shown in lighter shades. Values for active (red) and inactive (blue) equilibrium simulations are shown at the beginning of each microswitch plot. Values for transition trajectory (purple) follows. Mean values from equilibrium simulations are shown as red (active) and blue (inactive) horizontal lines.

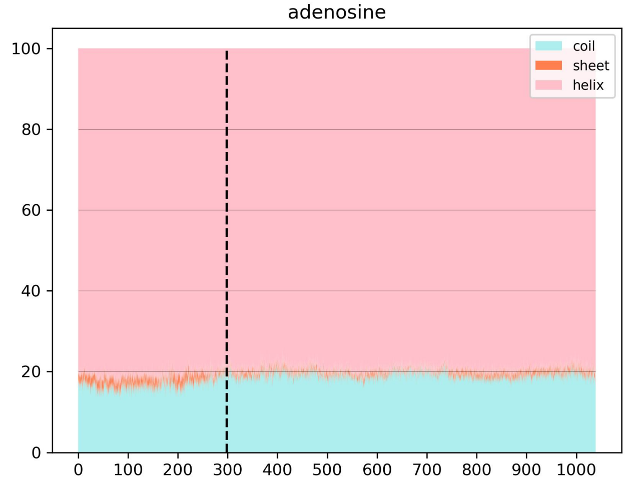


Figure 40: Secondary structure elements percentage for transition trajectory for adenosine system.

#### 1.25 orexin

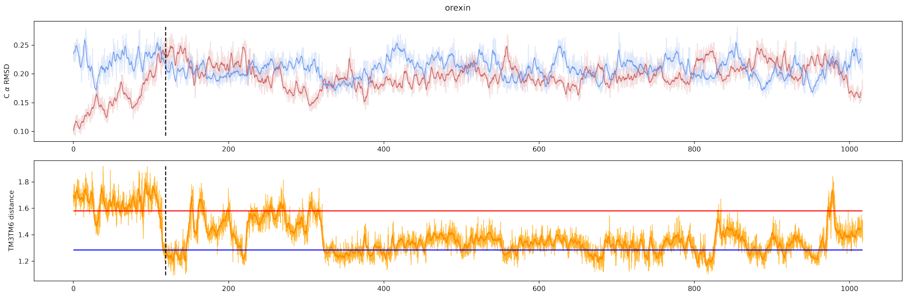


Figure 41: RMSD traces to active and inactive equilibrium ensembles (top) and TM3TM6 distance (bottom) for transition trajectory for orexin system. On both plots, running mean of size 25 (2.5 ns) shown in darker shades, and values for particular frames shown in lighter shades. Mean values of TM3TM6 distances from active (red) and inactive (blue) equilibrium simulations are shown as horizontal lines.

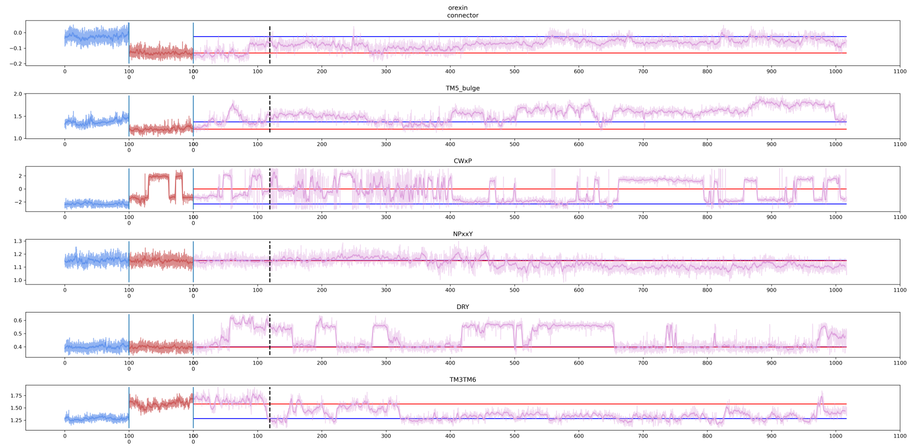


Figure 42: Selected microswitches changes in time for orexin system. On all plots, running mean of size 25 (2.5 ns, 1.25 ns for equilibria) shown in darker shades, and values for particular frames shown in lighter shades. Values for active (red) and inactive (blue) equilibrium simulations are shown at the beginning of each microswitch plot. Values for transition trajectory (purple) follows. Mean values from equilibrium simulations are shown as red (active) and blue (inactive) horizontal lines.

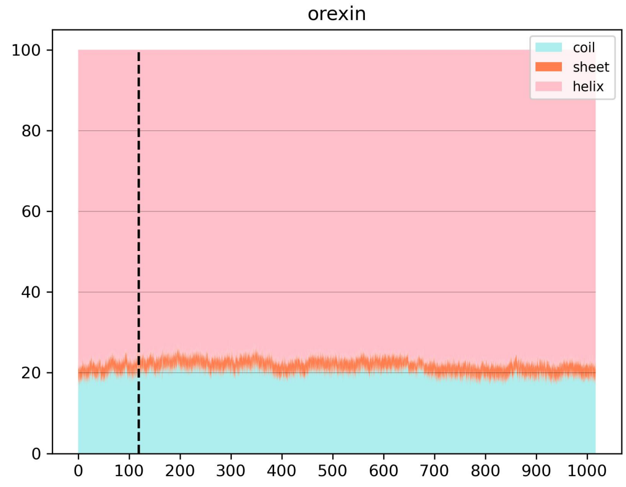


Figure 43: Secondary structure elements percentage for transition trajectory for orexin system.

#### 1.26 dopamine

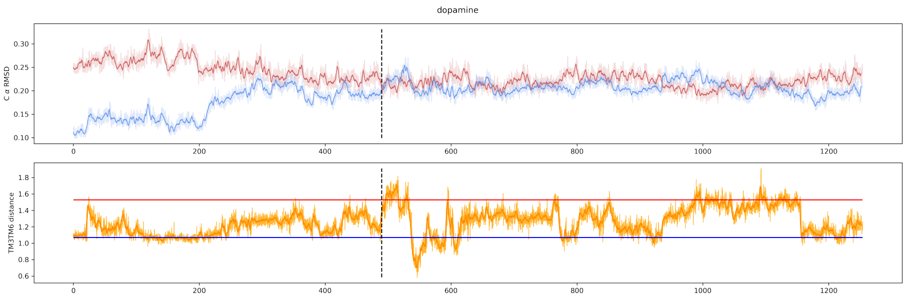


Figure 44: RMSD traces to active and inactive equilibrium ensembles (top) and TM3TM6 distance (bottom) for transition trajectory for dopamine system. On both plots, running mean of size 25 (2.5 ns) shown in darker shades, and values for particular frames shown in lighter shades. Mean values of TM3TM6 distances from active (red) and inactive (blue) equilibrium simulations are shown as horizontal lines.

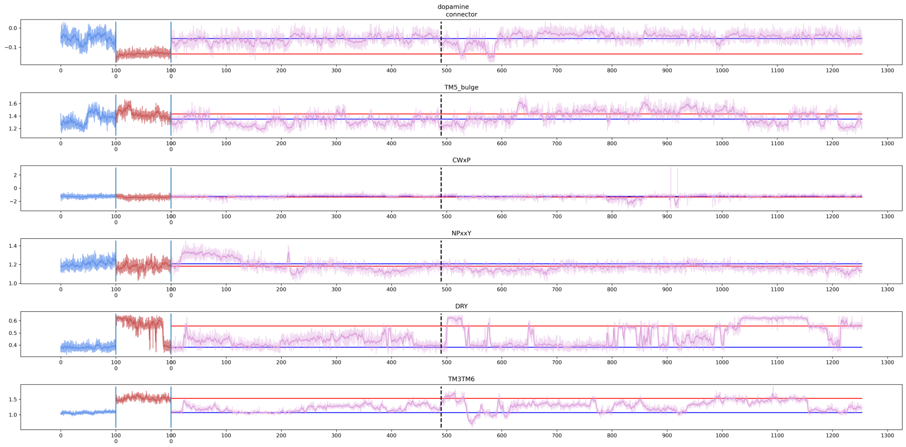


Figure 45: Selected microswitches changes in time for dopamine system. On all plots, running mean of size 25 (2.5 ns, 1.25 ns for equilibria) shown in darker shades, and values for particular frames shown in lighter shades. Values for active (red) and inactive (blue) equilibrium simulations are shown at the beginning of each microswitch plot. Values for transition trajectory (purple) follows. Mean values from equilibrium simulations are shown as red (active) and blue (inactive) horizontal lines.

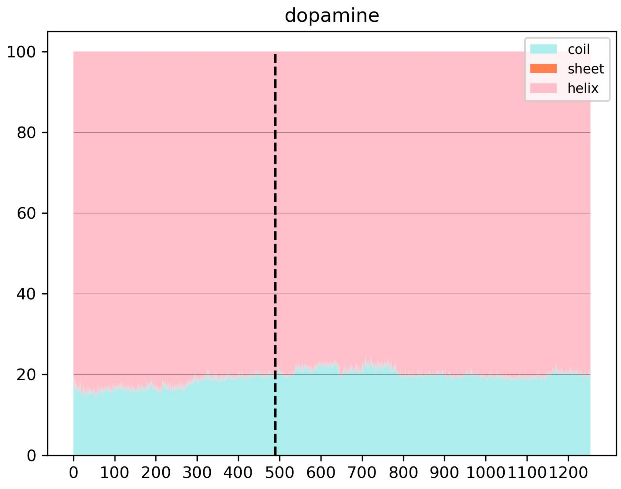


Figure 46: Secondary structure elements percentage for transition trajectory for dopamine system.

#### 1.27 angiotensin

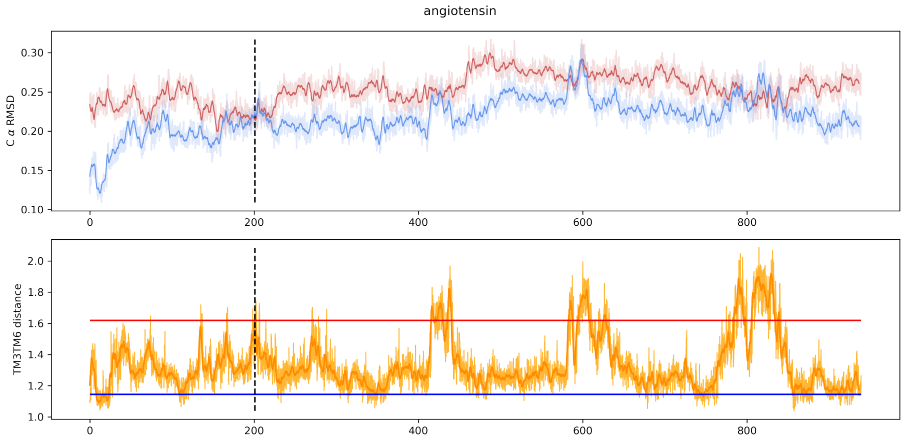


Figure 47: RMSD traces to active and inactive equilibrium ensembles (top) and TM3TM6 distance (bottom) for transition trajectory for angiotensin system. On both plots, running mean of size 25 (2.5 ns) shown in darker shades, and values for particular frames shown in lighter shades. Mean values of TM3TM6 distances from active (red) and inactive (blue) equilibrium simulations are shown as horizontal lines.

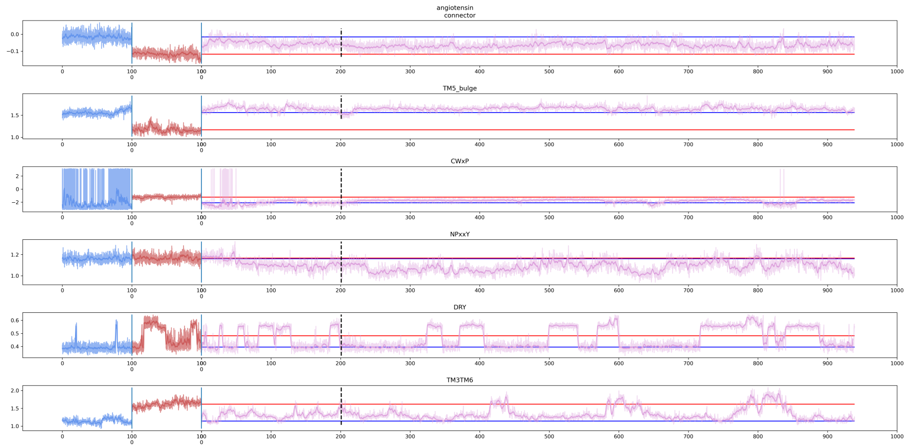


Figure 48: Selected microswitches changes in time for angiotensin system. On all plots, running mean of size 25 (2.5 ns, 1.25 ns for equilibria) shown in darker shades, and values for particular frames shown in lighter shades. Values for active (red) and inactive (blue) equilibrium simulations are shown at the beginning of each microswitch plot. Values for transition trajectory (purple) follows. Mean values from equilibrium simulations are shown as red (active) and blue (inactive) horizontal lines.

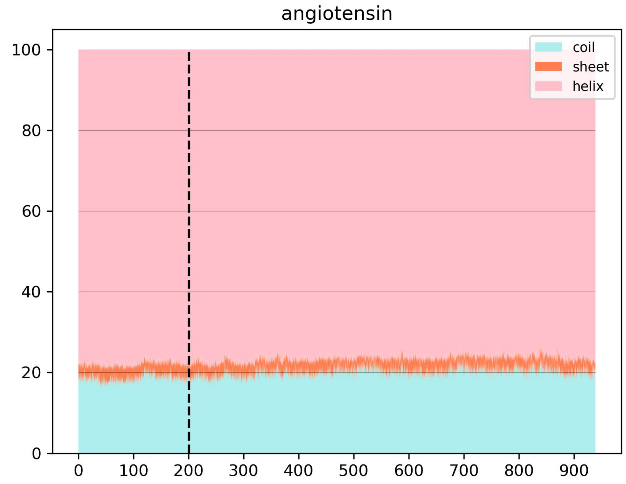


Figure 49: Secondary structure elements percentage for transition trajectory for angiotensin system.

#### 1.28 histamine

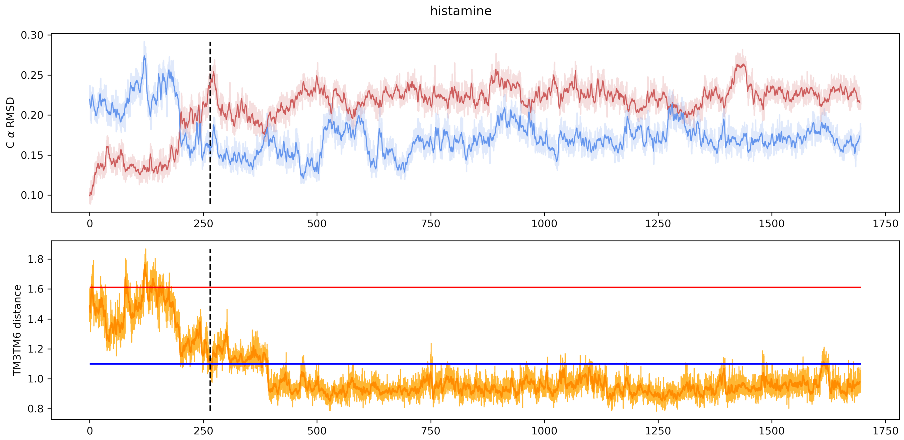


Figure 50: RMSD traces to active and inactive equilibrium ensembles (top) and TM3TM6 distance (bottom) for transition trajectory for histamine system. On both plots, running mean of size 25 (2.5 ns) shown in darker shades, and values for particular frames shown in lighter shades. Mean values of TM3TM6 distances from active (red) and inactive (blue) equilibrium simulations are shown as horizontal lines.

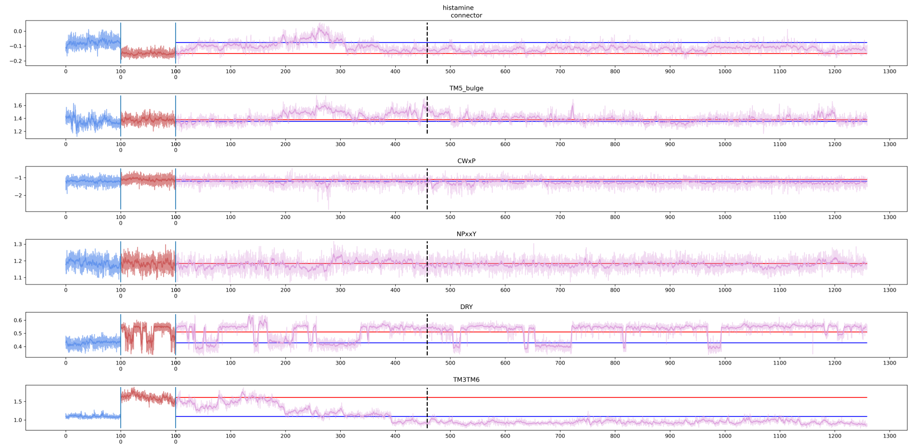


Figure 51: Selected microswitches changes in time for histamine system. On all plots, running mean of size 25 (2.5 ns, 1.25 ns for equilibria) shown in darker shades, and values for particular frames shown in lighter shades. Values for active (red) and inactive (blue) equilibrium simulations are shown at the beginning of each microswitch plot. Values for transition trajectory (purple) follows. Mean values from equilibrium simulations are shown as red (active) and blue (inactive) horizontal lines.

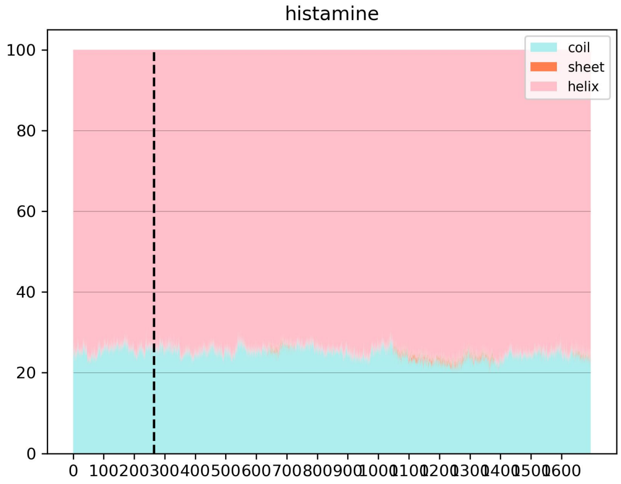


Figure 52: Secondary structure elements percentage for transition trajectory for histamine system.

#### 1.29 opsins

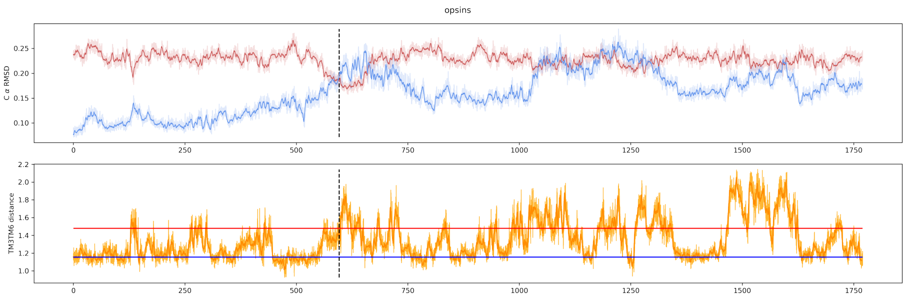


Figure 53: RMSD traces to active and inactive equilibrium ensembles (top) and TM3TM6 distance (bottom) for transition trajectory for opsins system. On both plots, running mean of size 25 (2.5 ns) shown in darker shades, and values for particular frames shown in lighter shades. Mean values of TM3TM6 distances from active (red) and inactive (blue) equilibrium simulations are shown as horizontal lines.

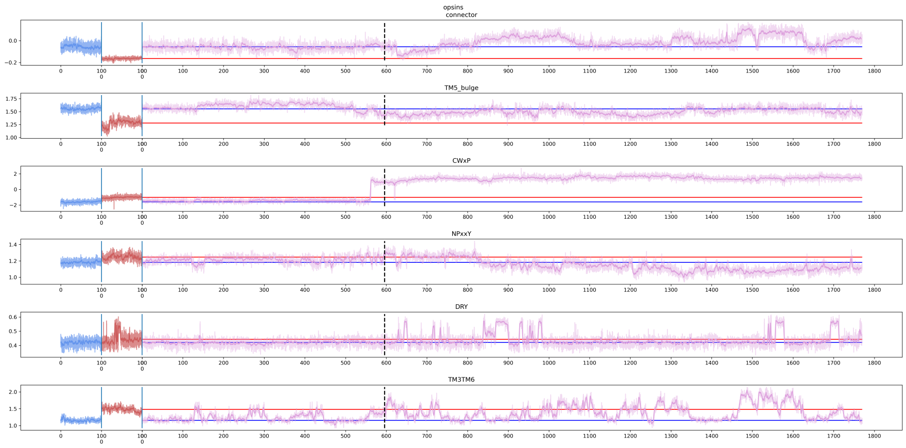


Figure 54: Selected microswitches changes in time for opsins system. On all plots, running mean of size 25 (2.5 ns, 1.25 ns for equilibria) shown in darker shades, and values for particular frames shown in lighter shades. Values for active (red) and inactive (blue) equilibrium simulations are shown at the beginning of each microswitch plot. Values for transition trajectory (purple) follows. Mean values from equilibrium simulations are shown as red (active) and blue (inactive) horizontal lines.

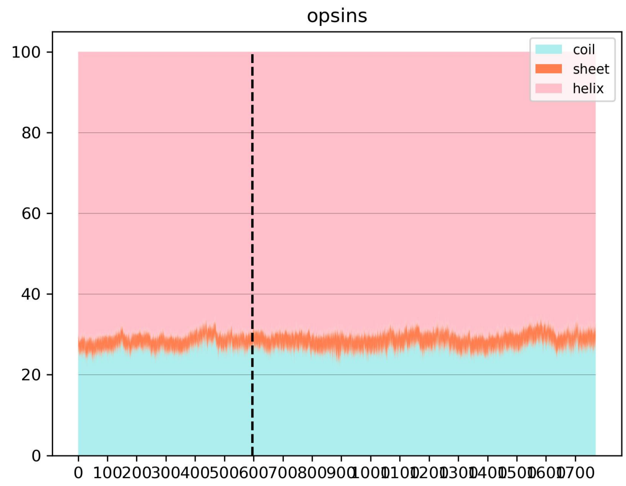


Figure 55: Secondary structure elements percentage for transition trajectory for opsins system.

#### 1.30 neurotensin

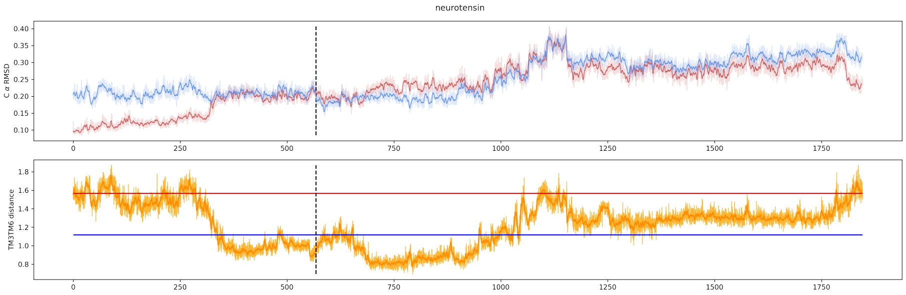


Figure 56: RMSD traces to active and inactive equilibrium ensembles (top) and TM3TM6 distance (bottom) for transition trajectory for neurotensin system. On both plots, running mean of size 25 (2.5 ns) shown in darker shades, and values for particular frames shown in lighter shades. Mean values of TM3TM6 distances from active (red) and inactive (blue) equilibrium simulations are shown as horizontal lines.

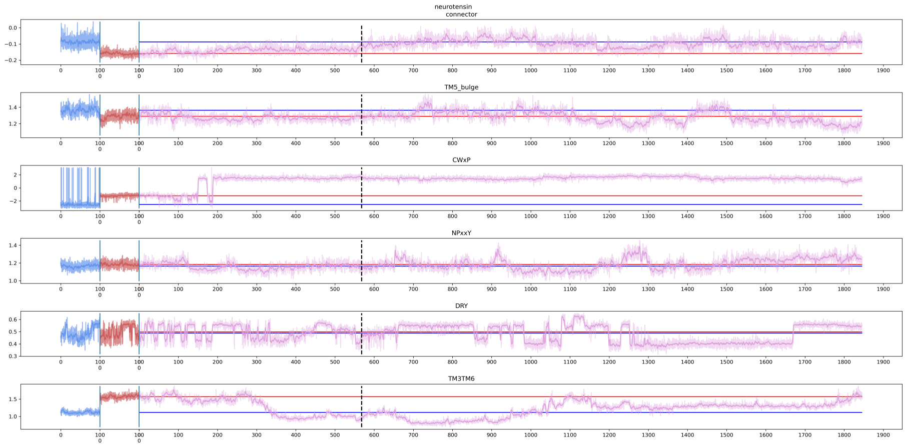


Figure 57: Selected microswitches changes in time for neurotensin system. On all plots, running mean of size 25 (2.5 ns, 1.25 ns for equilibria) shown in darker shades, and values for particular frames shown in lighter shades. Values for active (red) and inactive (blue) equilibrium simulations are shown at the beginning of each microswitch plot. Values for transition trajectory (purple) follows. Mean values from equilibrium simulations are shown as red (active) and blue (inactive) horizontal lines.

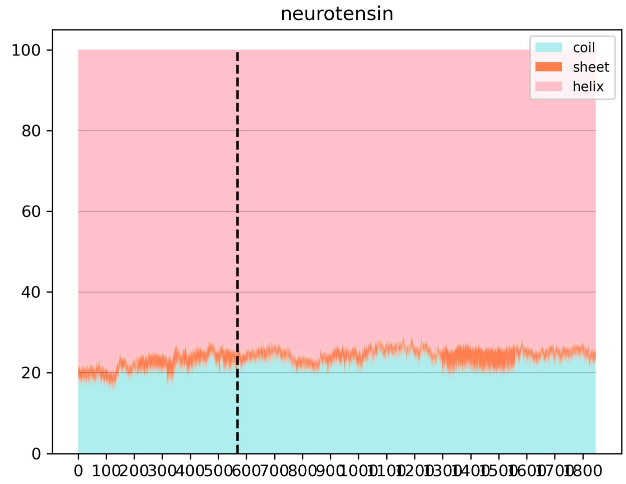


Figure 58: Secondary structure elements percentage for transition trajectory for neurotensin system.

#### 1.31 adrb2

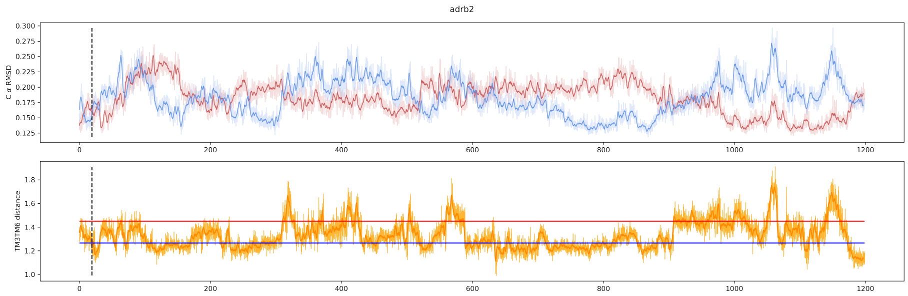


Figure 59: RMSD traces to active and inactive equilibrium ensembles (top) and TM3TM6 distance (bottom) for transition trajectory for adrb2 system. On both plots, running mean of size 25 (2.5 ns) shown in darker shades, and values for particular frames shown in lighter shades. Mean values of TM3TM6 distances from active (red) and inactive (blue) equilibrium simulations are shown as horizontal lines.

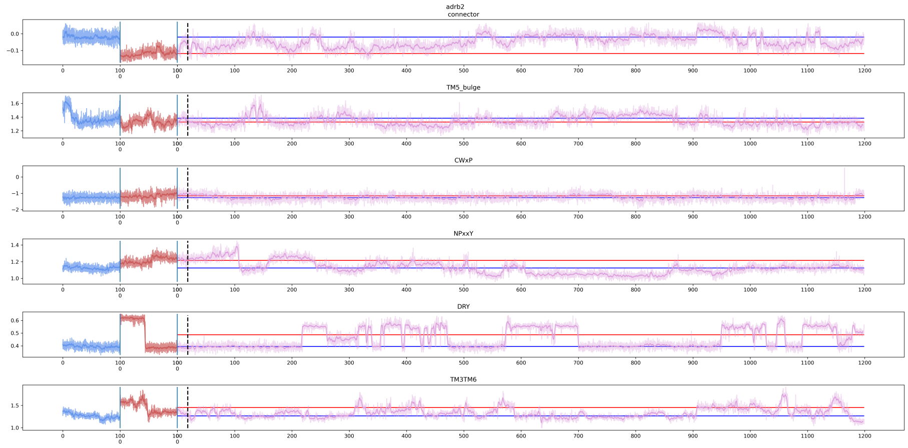


Figure 60: Selected microswitches changes in time for adrb2 system. On all plots, running mean of size 25 (2.5 ns, 1.25 ns for equilibria) shown in darker shades, and values for particular frames shown in lighter shades. Values for active (red) and inactive (blue) equilibrium simulations are shown at the beginning of each microswitch plot. Values for transition trajectory (purple) follows. Mean values from equilibrium simulations are shown as red (active) and blue (inactive) horizontal lines.

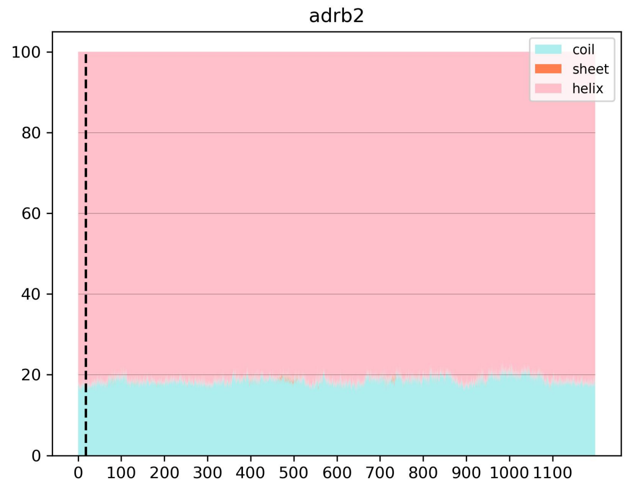


Figure 61: Secondary structure elements percentage for transition trajectory for adrb2 system.

#### 1.32 GPR183

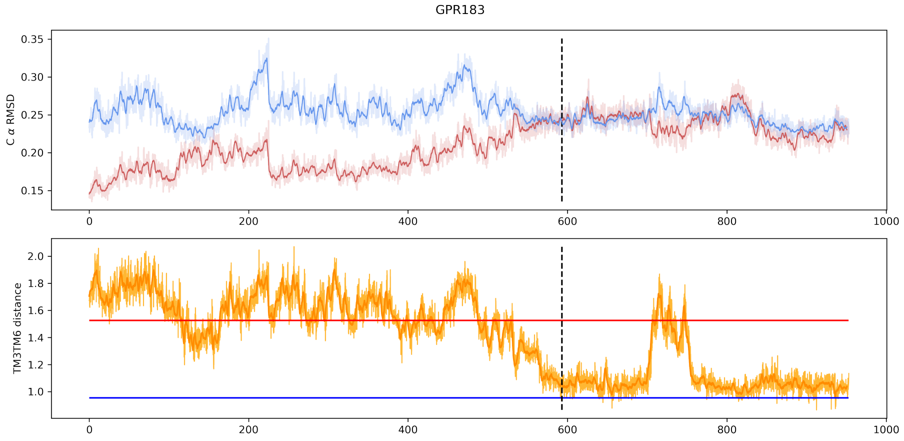


Figure 62: RMSD traces to active and inactive equilibrium ensembles (top) and TM3TM6 distance (bottom) for transition trajectory for GPR183 system. On both plots, running mean of size 25 (2.5 ns) shown in darker shades, and values for particular frames shown in lighter shades. Mean values of TM3TM6 distances from active (red) and inactive (blue) equilibrium simulations are shown as horizontal lines.

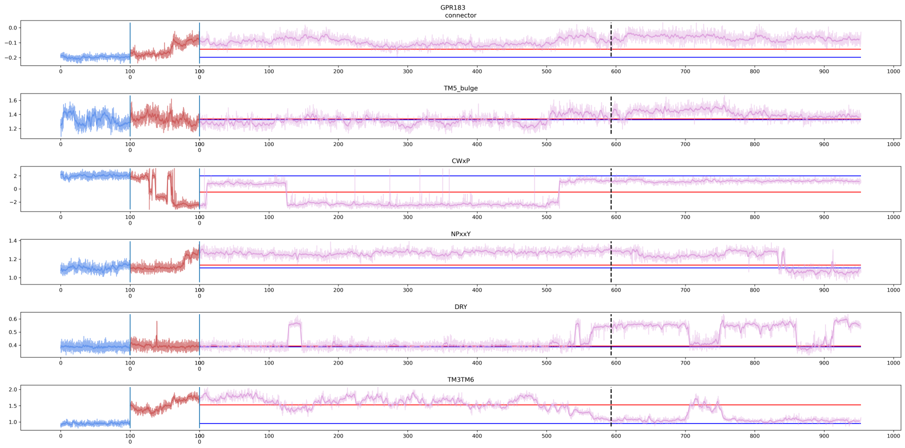


Figure 63: Selected microswitches changes in time for GPR183 system. On all plots, running mean of size 25 (2.5 ns, 1.25 ns for equilibria) shown in darker shades, and values for particular frames shown in lighter shades. Values for active (red) and inactive (blue) equilibrium simulations are shown at the beginning of each microswitch plot. Values for transition trajectory (purple) follows. Mean values from equilibrium simulations are shown as red (active) and blue (inactive) horizontal lines.

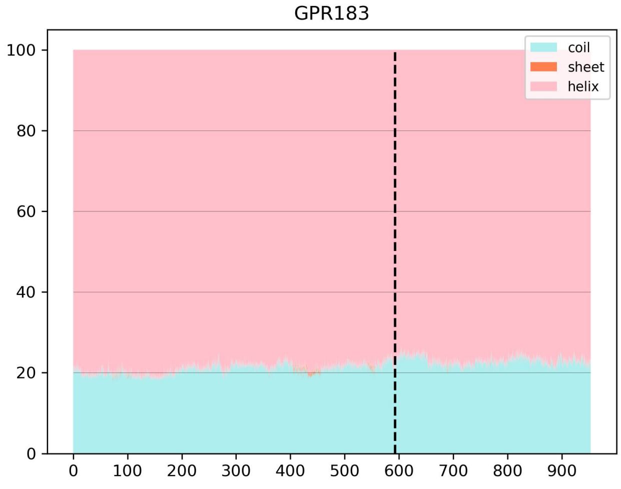


Figure 64: Secondary structure elements percentage for transition trajectory for GPR183 system.

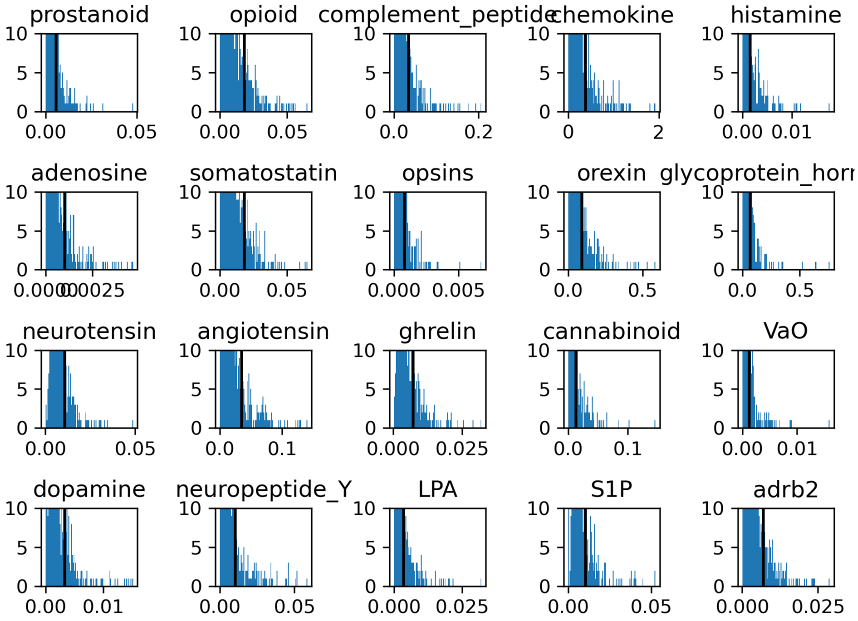


Figure 65: Histograms of feature importances for latent dimension 1 for all the systems. Vertical lines indicate top 100.

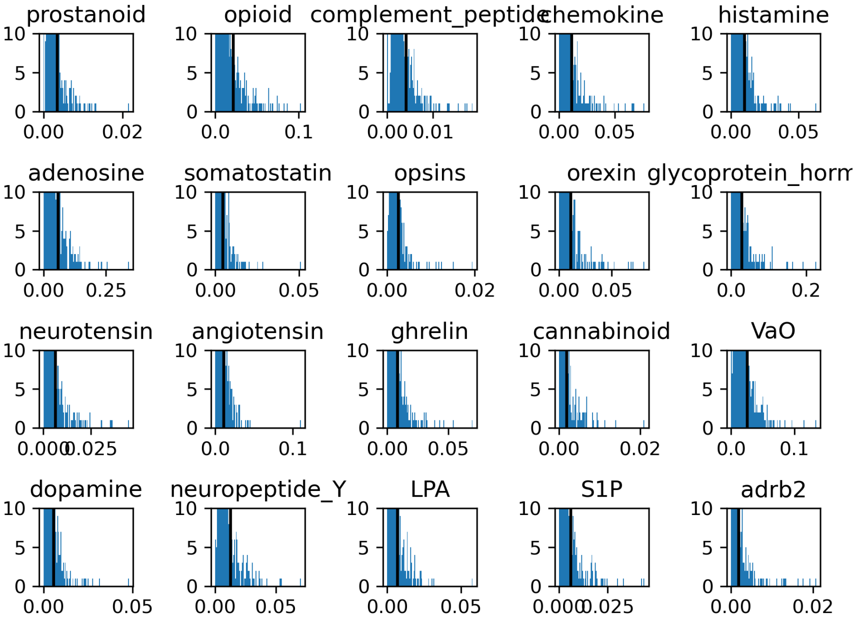


Figure 66: Histograms of feature importances for latent dimension 2 for all the systems. Vertical lines indicate top 100.

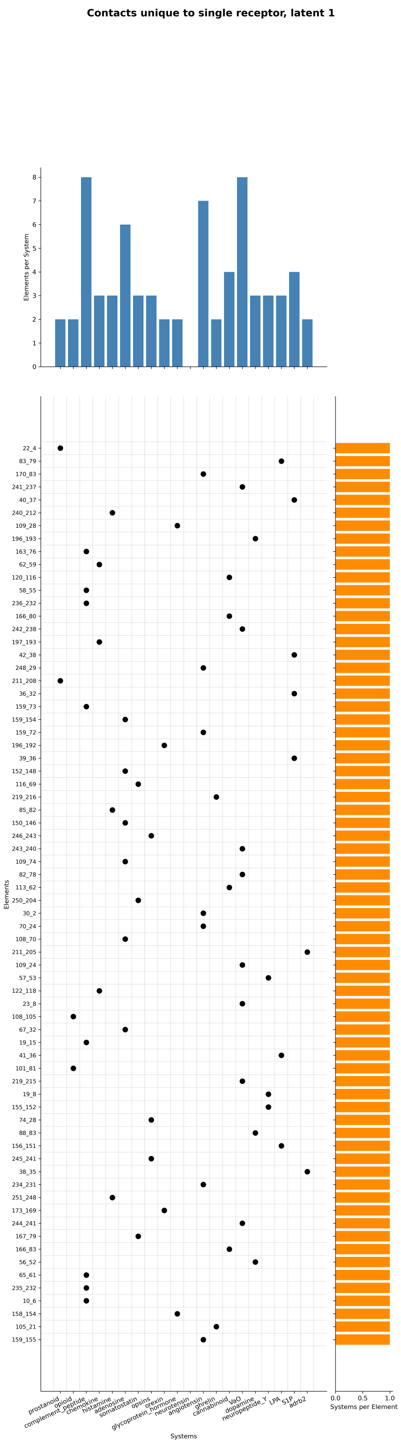


Figure 67: Residue pairs included in top 100 most important features only for a single receptor, for latent dimension 1.

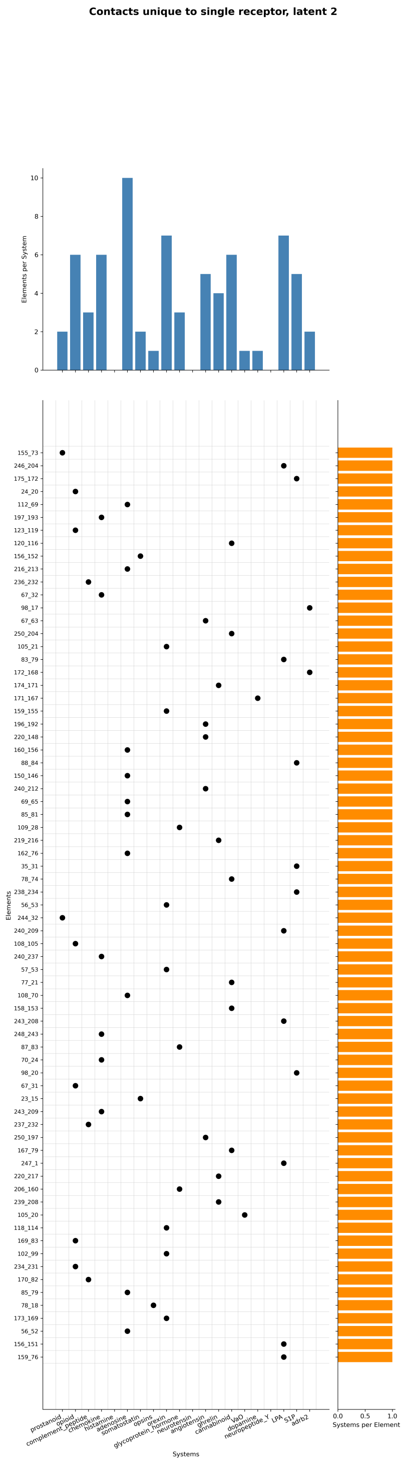


Figure 68: Residue pairs included in top 100 most important features only for a single receptor, for latent dimension 2.

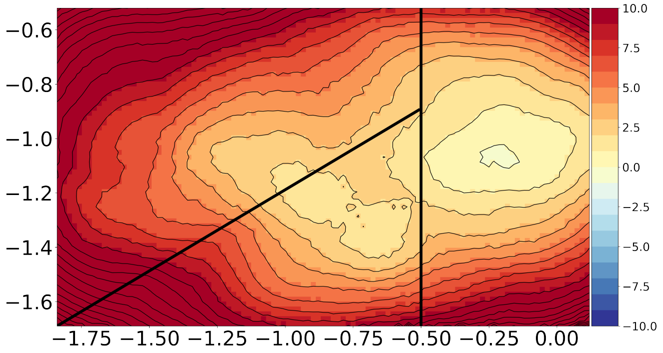


Figure 69: Boundaries of the states. For state proportions, unnormalized probability has been computed based on the whole Free Energy Surface, each region has been summed over and proportions between them have been used to determine ligand activity in the Fig. S7.

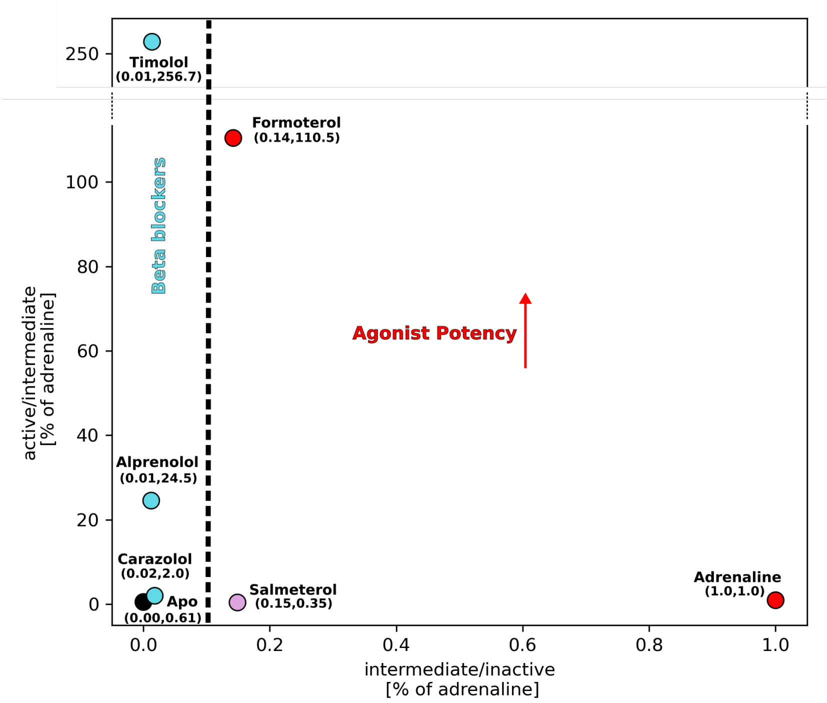


Figure 70: Relationship between ligand modality and state proportions. Beta blockers have low proportions between intermediate and inactive states( 0.01). If a ligand has higher proportions( 0.14), it is an agonist. For an agonist, proportions between active and intermediate states indicate potency: for partial agonist salmeterol it is 0.35, and for potent synthetic agonist formoterol it is 110.5. Values were normalized to adrenaline values.

##### 1.32.1 Formoterol

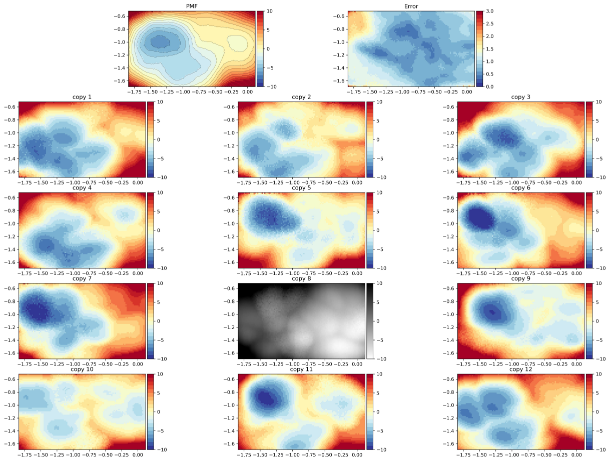


Figure 71: Free energy surfaces of $\beta$2-adrenergic receptor activation bound with synthetic agonist formoterol. Mean FES shown on the top left, and jackknife error shown on the top right. Final FES from all individual copies shown below them. All values in kcal/mol. An outlier copy excluded from final results shown in gray.

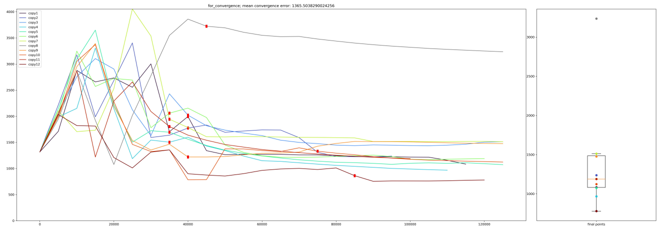

Figure 72: Convergence of FES simulations with formoterol for each individual copy. We visualized distances between profiles in time, with 1 ns spacing. An outlier copy excluded from final results shown in gray.

##### 1.32.2 Adrenaline

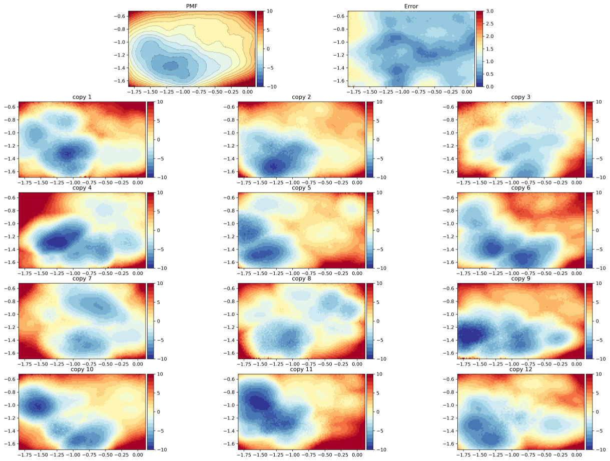

Figure 73: Free energy surfaces of $\beta$2-adrenergic receptor activation bound with orthosteric agonist adrenaline. Mean FES shown on the top left, and jackknife error shown on the top right. Final FES from all individual copies shown below them. All values in kcal/mol.

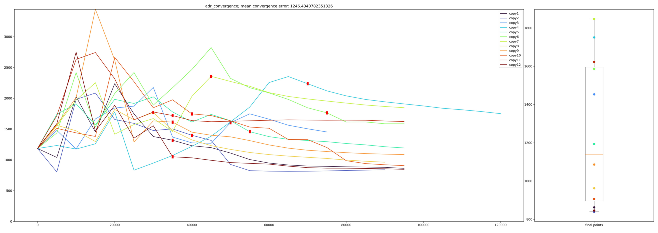

Figure 74: Convergence of FES simulations with adrenaline for each individual copy. Distances between profiles in time are shown, with 1 ns spacing.

##### 1.32.3 Salmeterol

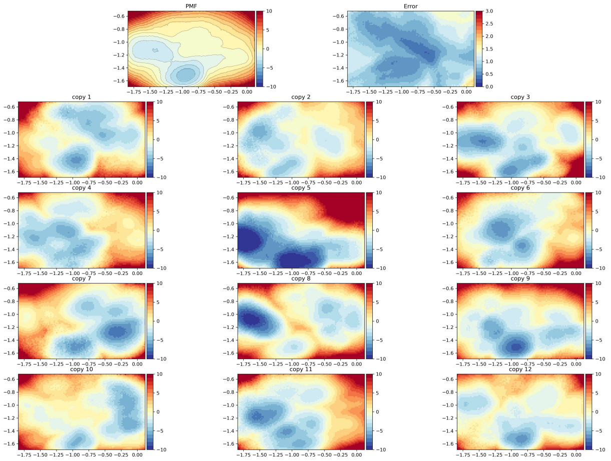


Figure 75: Free energy surfaces of $\beta$2-adrenergic receptor activation bound with partial agonist salmeterol. Mean FES shown on the top left, and jackknife error shown on the top right. Final FES from all individual copies shown below them. All values in kcal/mol.

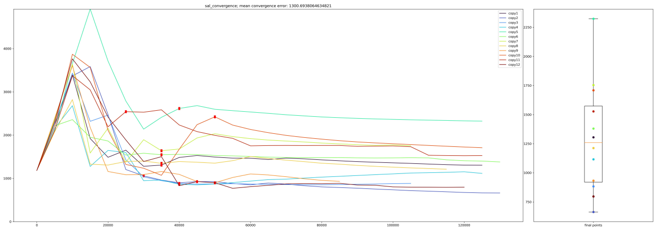

Figure 76: Convergence of FES simulations with salmeterol for each individual copy. Distances between profiles in time are shown, with 1 ns spacing.

##### 1.32.4 Apo

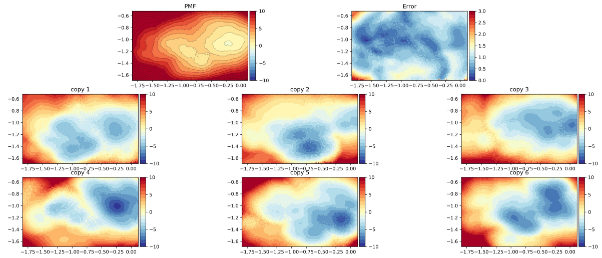


Figure 77: Free energy surfaces of $\beta$2-adrenergic receptor activation in apo conditions. Mean FES shown on the top left, and jackknife error shown on the top right. Final FES from all individual copies shown below them. All values in kcal/mol.

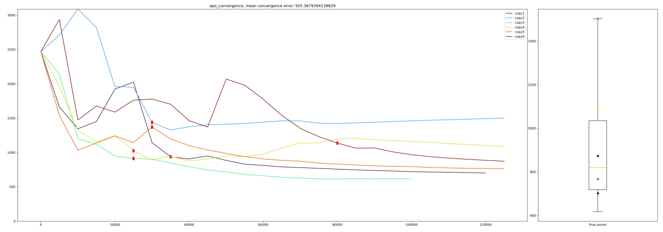

Figure 78: Convergence of FES simulations in apo condition for each individual copy. Distances between profiles in time are shown, with 1 ns spacing.

##### 1.32.5 Carazolol

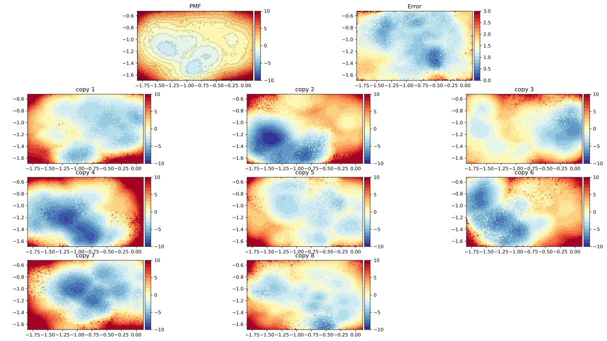


Figure 79: Free energy surfaces of $\beta$2-adrenergic receptor activation bound with beta blocker carazolol. Mean FES shown on the top left, and jackknife error shown on the top right. Final FES from all individual copies shown below them. All values in kcal/mol. This profile looks more rough than others because of more extensive sampling between AWH FES updates than with other ligands (nstsample=1000 instead of 100)

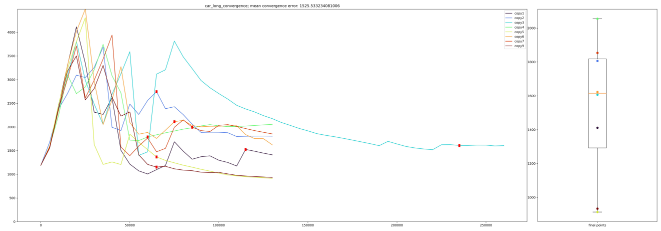

Figure 80: Convergence of FES simulations with carazolol for each individual copy. Distances between profiles in time are shown, with 1 ns spacing.

##### 1.32.6 Timolol

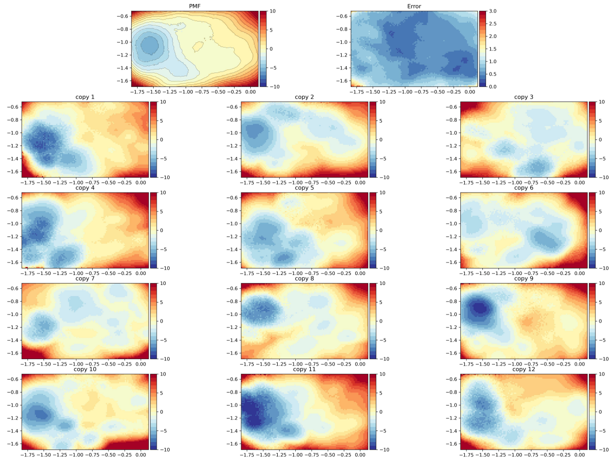


Figure 81: Free energy surfaces of $\beta$2-adrenergic receptor activation bound with beta blocker timolol. Mean FES shown on the top left, and jackknife error shown on the top right. Final FES from all individual copies shown below them. All values in kcal/mol.

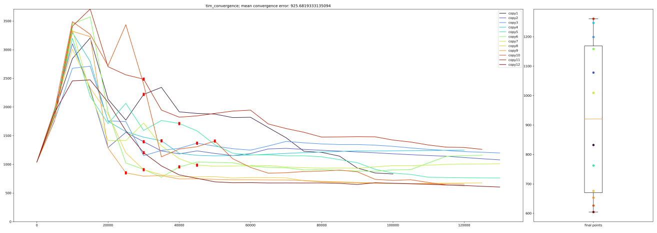

Figure 82: Convergence of FES simulations with timolol for each individual copy. Distances between profiles in time are shown, with 1 ns spacing.

##### 1.32.7 Alprenolol

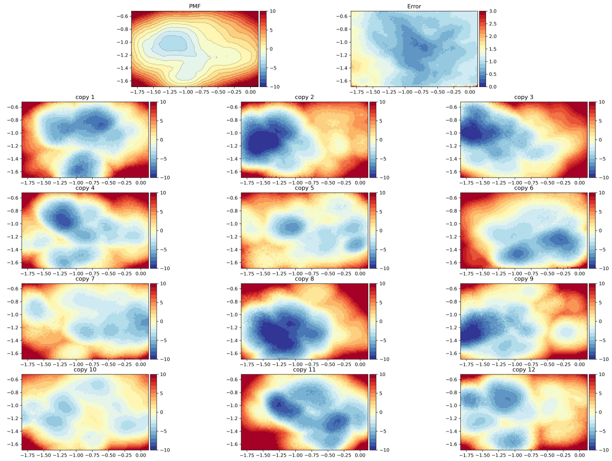


Figure 83: Free energy surfaces of $\beta$2-adrenergic receptor activation bound with beta blocker alprenolol. Mean FES shown on the top left, and jackknife error shown on the top right. Final FES from all individual copies shown below them. All values in kcal/mol.

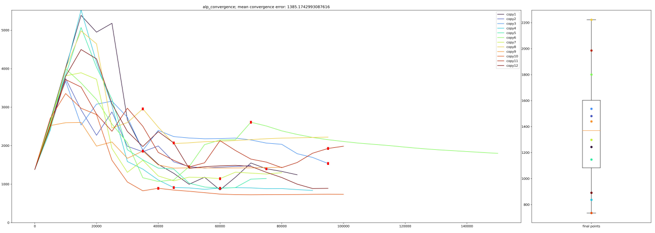

Figure 84: Convergence of FES simulations with alprenolol for each individual copy. Distances between profiles in time are shown, with 1 ns spacing.

[5] Pfam: The protein families database in 2021 | Nucleic Acids Research | Oxford Academic.

[6] UniProt: the Universal Protein Knowledgebase in 2025 | Nucleic Acids Research | Oxford Academic.

[7] Comparative Protein Structure Modeling Using MODELLER - Webb - 2016 - Current Protocols in Bioinformatics - Wiley Online Library.

[15] Improved Treatment of Ligands and Coupling Effects in Empirical Calculation and Rationalization of pKa Values | Journal of Chemical Theory and Computation.

[16] H. J. C. Berendsen, D. van der Spoel, and R. van Drunen. GROMACS: A message-passing parallel molecular dynamics implementation. *Computer Physics Communications*, 91(1):43–56, September 1995.

[17] David Van Der Spoel, Erik Lindahl, Berk Hess, Gerrit Groenhof, Alan E. Mark, and Herman J. C. Berendsen. GROMACS: Fast, flexible, and free. *Journal of Computational Chemistry*, 26(16):1701–1718, December 2005.

[18] Erik Lindahl, Berk Hess, and David van der Spoel. GROMACS 3.0: a package for molecular simulation and trajectory analysis. *Molecular modeling annual*, 7(8):306–317, August 2001.

[19] Berk Hess, Carsten Kutzner, David van der Spoel, and Erik Lindahl. GROMACS 4: Algorithms for Highly Efficient, Load-Balanced, and Scalable Molecular Simulation. *Journal of Chemical Theory and Computation*, 4(3):435–447, March 2008.

[20] GROMACS 4.5: a high-throughput and highly parallel open source molecular simulation toolkit | Bioinformatics | Oxford Academic.

[21] Mark James Abraham, Teemu Murtola, Roland Schulz, Szilárd Páll, Jeremy C. Smith, Berk Hess, and Erik Lindahl. GROMACS: High performance molecular simulations through multi-level parallelism from laptops to supercomputers. *SoftwareX*, 1-2:19–25, September 2015.

[22] Páll Szilárd, Mark James Abraham, Carsten Kutzner, Berk Hess, and Erik Lindahl. Tackling Exascale Software Challenges in Molecular Dynamics Simulations with GROMACS. volume 8759, pages 3–27. 2015. arXiv:1506.00716 [cs.CE].

[23] B. R. Brooks, C. L. Brooks III, A. D. Mackerell Jr., L. Nilsson, R. J. Petrella, B. Roux, Y. Won, G. Archontis, C. Bartels, S. Boresch, A. Caflisch, L. Caves, Q. Cui, A. R. Dinner, M. Feig, S. Fischer, J. Gao, M. Hodoscek, W. Im, K. Kuczera, T. Lazaridis, J. Ma, V. Ovchinnikov, E. Paci, R. W. Pastor, C. B. Post, J. Z. Pu, M. Schaefer, B. Tidor, R. M. Venable, H. L. Woodcock, X. Wu, W. Yang, D. M. York, and M. Karplus. CHARMM: The biomolecular simulation program. *Journal of Computational Chemistry*, 30(10):1545–1614, 2009. _eprint: https://onlinelibrary.wiley.com/doi/pdf/10.1002/jcc.21287.

[24] Pressure control using stochastic cell rescaling | The Journal of Chemical Physics | AIP Publishing.

[41] Shen-Ming Huang, Meng-Yao Xiong, Lei Liu, Jianqiang Mu, Ming-Wei Wang, Ying-Li Jia, Kui Cai, Lu Tie, Chao Zhang, Sheng Cao, Xin Wen, Jia-Le Wang, Sheng-Chao Guo, Yu Li, Chang-Xiu Qu, Qing-Tao He, Bo-Yang Cai, Chenyang Xue, Shiyi Gan, Yihe Xie, Xin Cong, Zhao Yang, Wei Kong, Shuo Li, Zijian Li, Peng Xiao, Fan Yang, Xiao Yu, You-Fei Guan, Xiaoyan Zhang, Zhongmin Liu, Bao-Xue Yang, Yang Du, and Jin-Peng Sun. Single hormone or synthetic agonist induces Gs/Gi coupling selectivity of EP receptors via distinct binding modes and propagating paths. *Proceedings of the National Academy of Sciences*, 120(30):e2216329120, July 2023.

[42] Yosuke Toyoda, Kazushi Morimoto, Ryoji Suno, Shoichiro Horita, Keitaro Yamashita, Kunio Hirata, Yusuke Sekiguchi, Satoshi Yasuda, Mitsunori Shiroishi, Tomoko Shimizu, Yuji Urushibata, Yuta Kajiwara, Tomoaki Inazumi, Yunhon Hotta, Hidetsugu Asada, Takanori Nakane, Yuki Shimura, Tomoya Nakagita, Kyoshiro Tsuge, Suguru Yoshida, Tomoko Kuribara, Takamitsu Hosoya, Yukihiko Sugimoto, Norimichi Nomura, Miwa Sato, Takatsugu Hirokawa, Masahiro Kinoshita, Takeshi Murata, Kiyoshi Takayama, Masaki Yamamoto, Shuh Narumiya, So Iwata, and Takuya Kobayashi. Ligand binding to human prostaglandin E receptor EP4 at the lipid-bilayer interface. *Nature Chemical Biology*, 15(1):18–26, January 2019.

[49] Receptor-specific recognition of NPY peptides revealed by structures of NPY receptors | Science Advances.

[50] Zhenlin Yang, Shuo Han, Max Keller, Anette Kaiser, Brian J. Bender, Mathias Bosse, Kerstin Burkert, Lisa M. Kögler, David Wifling, Guenther Bernhardt, Nicole Plank, Timo Littmann, Peter Schmidt, Cuiying Yi, Beibei Li, Sheng Ye, Rongguang Zhang, Bo Xu, Dan Larhammar, Raymond C. Stevens, Daniel Huster, Jens Meiler, Qiang Zhao, Annette G. Beck-Sickinger, Armin Buschauer, and Beili Wu. Structural basis of ligand binding modes at the neuropeptide Y Y1 receptor. *Nature*, 556(7702):520–524, April 2018.

[51] Jiao Qin, Ye Cai, Zheng Xu, Qianqian Ming, Su-Yu Ji, Chao Wu, Huibing Zhang, Chunyou Mao, Dan-Dan Shen, Kunio Hirata, Yanbin Ma, Wei Yan, Yan Zhang, and Zhenhua Shao. Molecular mechanism of agonism and inverse agonism in ghrelin receptor. *Nature Communications*, 13(1):300, January 2022.

[52] Yuki Shiimura, Shoichiro Horita, Akie Hamamoto, Hidetsugu Asada, Kunio Hirata, Misuzu Tanaka, Kenji Mori, Tomoko Uemura, Takuya Kobayashi, So Iwata, and Masayasu Kojima. Structure of an antagonist-bound ghrelin receptor reveals possible ghrelin recognition mode. *Nature Communications*, 11(1):4160, August 2020.

[53] Shian Liu, Navid Paknejad, Lan Zhu, Yasuyuki Kihara, Manisha Ray, Jerold Chun, Wei Liu, Richard K. Hite, and Xin-Yun Huang. Differential activation mechanisms of lipid GPCRs by lysophosphatidic acid and sphingosine 1-phosphate. *Nature Communications*, 13(1):731, February 2022.

[54] Crystal Structure of a Lipid G Protein–Coupled Receptor | Science.

[55] Jill E. Chrencik, Christopher B. Roth, Masahiko Terakado, Haruto Kurata, Rie Omi, Yasuyuki Kihara, Dora Warshaviak, Shinji Nakade, Guillermo Asmar-Rovira, Mauro Mileni, Hirotaka Mizuno, Mark T. Griffith, Caroline Rodgers, Gye Won Han, Jeffrey Velasquez, Jerold Chun, Raymond C. Stevens, and Michael A. Hanson. Crystal Structure of Antagonist Bound Human Lysophosphatidic Acid Receptor 1. *Cell*, 161(7):1633–1643, June 2015.

[56] Bryan Faust, Christian B. Billesbølle, Carl-Mikael Suomivuori, Isha Singh, Kaihua Zhang, Nicholas Hoppe, Antonio F. M. Pinto, Jolene K. Diedrich, Yagmur Muftuoglu, Mariusz W. Szkudlinski, Alan Saghatelian, Ron O.

Dror, Yifan Cheng, and Aashish Manglik. Autoantibody mimicry of hormone action at the thyrotropin receptor. *Nature*, 609(7928):846–853, September 2022.

[57] Justin G. Meyerowitz, Michael J. Robertson, Ximena Barros-Álvarez, Ouliana Panova, Robert M. Nwokonko, Yang Gao, and Georgios Skiniotis. The oxytocin signaling complex reveals a molecular switch for cation dependence. *Nature Structural & Molecular Biology*, 29(3):274–281, March 2022.

[58] Yann Waltenspühl, Jendrik Schöppe, Janosch Ehrenmann, Lutz Kummer, and Andreas Plückthun. Crystal structure of the human oxytocin receptor. *Science Advances*, 6(29):eabb5419, July 2020.

[59] Christopher J. Draper-Joyce, Rebecca Bhola, Jinan Wang, Apurba Bhattarai, Anh T. N. Nguyen, India Cowie-Kent, Kelly O'Sullivan, Ling Yeong Chia, Hariprasad Venugopal, Celine Valant, David M. Thal, Denise Wootten, Nicolas Panel, Jens Carlsson, Macdonald J. Christie, Paul J. White, Peter Scammells, Lauren T. May, Patrick M. Sexton, Radostin Danev, Yinglong Miao, Alisa Glukhova, Wendy L. Imlach, and Arthur Christopoulos. Positive allosteric mechanisms of adenosine A1 receptor-mediated analgesia. *Nature*, 597(7877):571–576, September 2021.

[60] Structure of the Adenosine A1 Receptor Reveals the Basis for Subtype Selectivity: Cell.

[61] Chuan Hong, Noel J. Byrne, Beata Zamlynny, Srivanya Tummala, Li Xiao, Jennifer M. Shipman, Andrea T. Partridge, Christina Minnick, Michael J. Breslin, Michael T. Rudd, Shawn J. Stachel, Vanessa L. Rada, Jeffrey C. Kern, Kira A. Armacost, Scott A. Hollingsworth, Julie A. O'Brien, Dawn L. Hall, Terrence P. McDonald, Corey Strickland, Alexei Brooun, Stephen M. Soisson, and Kaspar Hollenstein. Structures of active-state orexin receptor 2 rationalize peptide and small-molecule agonist recognition and receptor activation. *Nature Communications*, 12(1):815, February 2021.

[62] Crystal Structures of Human Orexin 2 Receptor Bound to the Subtype-Selective Antagonist EMPA: Structure.

[63] Structural insights into the human D1 and D2 dopamine receptor signaling complexes: Cell.

[64] Sheng Wang, Tao Che, Anat Levit, Brian K. Shoichet, Daniel Wacker, and Bryan L. Roth. Structure of the D2 dopamine receptor bound to the atypical antipsychotic drug risperidone. *Nature*, 555(7695):269–273, March 2018.

[65] Dongqi Zhang, Yongfeng Liu, Saheem A. Zaidi, Lingyi Xu, Yuting Zhan, Anqi Chen, Jiangtao Guo, Xi-Ping Huang, Bryan L. Roth, Vsevolod Katritch, Vadim Cherezov, and Haitao Zhang. Structural insights into an-

giotensin receptor signaling modulation by balanced and biased agonists. *The EMBO Journal*, 42(11):EMBJ2022112940, April 2023.

[66] Structural Basis for Ligand Recognition and Functional Selectivity at Angiotensin Receptor* - Journal of Biological Chemistry.

[67] Ruixue Xia, Na Wang, Zhenmei Xu, Yang Lu, Jing Song, Anqi Zhang, Changyou Guo, and Yuanzheng He. Cryo-EM structure of the human histamine H1 receptor/Gq complex. *Nature Communications*, 12(1):2086, April 2021.

[68] Tatsuro Shimamura, Mitsunori Shiroishi, Simone Weyand, Hirokazu Tsujimoto, Graeme Winter, Vsevolod Katritch, Ruben Abagyan, Vadim Cherezov, Wei Liu, Gye Won Han, Takuya Kobayashi, Raymond C. Stevens, and So Iwata. Structure of the human histamine H1 receptor complex with doxepin. *Nature*, 475(7354):65–70, July 2011.

[69] Yang Gao, Hongli Hu, Sekar Ramachandran, Jon W. Erickson, Richard A. Cerione, and Georgios Skiniotis. Structures of the Rhodopsin-Transducin Complex: Insights into G-Protein Activation. *Molecular Cell*, 75(4):781–790.e3, August 2019.

[70] Thomas Gruhl, Tobias Weinert, Matthew J. Rodrigues, Christopher J. Milne, Giorgia Ortolani, Karol Nass, Eriko Nango, Saumik Sen, Philip J. M. Johnson, Claudio Cirelli, Antonia Furrer, Sandra Mous, Petr Skopintsev, Daniel James, Florian Dworkowski, Petra Båth, Demet Kekilli, Dmitry Ozerov, Rie Tanaka, Hannah Glover, Camila Bacellar, Steffen Brünle, Cecilia M. Casadei, Azeglio D. Diethelm, Dardan Gashi, Guillaume Gotthard, Ramon Guixà-González, Yasumasa Joti, Victoria Kabanova, Gregor Knopp, Elena Lesca, Pikyee Ma, Isabelle Martiel, Jonas Mühle, Shigeki Owada, Filip Pamula, Daniel Sarabi, Oliver Tejero, Ching-Ju Tsai, Niranjan Varma, Anna Wach, Sébastien Boutet, Kensuke Tono, Przemyslaw Nogly, Xavier Deupi, So Iwata, Richard Neutze, Jörg Standfuss, Gebhard Schertler, and Valerie Panneels. Ultrafast structural changes direct the first molecular events of vision. *Nature*, 615(7954):939–944, March 2023.

[71] Hideaki E. Kato, Yan Zhang, Hongli Hu, Carl-Mikael Suomivuori, Francois Marie Ngako Kadji, Junken Aoki, Kaavya Krishna Kumar, Rasmus Fonseca, Daniel Hilger, Weijiao Huang, Naomi R. Latorraca, Asuka Inoue, Ron O. Dror, Brian K. Kobilka, and Georgios Skiniotis. Conformational transitions of a neurotensin receptor 1–Gi1 complex. *Nature*, 572(7767):80–85, August 2019.

[72] Structure of signaling-competent neurotensin receptor 1 obtained by directed evolution in Escherichia coli | PNAS.

[73] High-Resolution Crystal Structure of an Engineered Human 2-Adrenergic G Protein–Coupled Receptor | Science.

[74] Hongwen Chen, Weijiao Huang, and Xiaochun Li. Structures of oxysterol sensor EBI2/GPR183, a key regulator of the immune response. *Structure*, 30(7):1016–1024.e5, July 2022.

[75] Peter J. A. Cock, Tiago Antao, Jeffrey T. Chang, Brad A. Chapman, Cymon J. Cox, Andrew Dalke, Iddo Friedberg, Thomas Hamelryck, Frank Kauff, Bartek Wilczynski, and Michiel J. L. de Hoon. Biopython: freely available Python tools for computational molecular biology and bioinformatics. *Bioinformatics*, 25(11):1422–1423, June 2009.

[76] Ivica Letunic and Peer Bork. Interactive Tree of Life (iTOL) v6: recent updates to the phylogenetic tree display and annotation tool. *Nucleic Acids Research*, 52(W1):W78–W82, July 2024.
